# Metabolic alterations in the absence of a detectable neuromuscular phenotype in novel genomically humanised *SOD1^A4V^* mice

**DOI:** 10.1101/2025.05.01.650597

**Authors:** David Thompson, Chloe Williams, Andrew P. Tosolini, Jonathan Gilthorpe, Giampietro Schiavo, Elizabeth M.C. Fisher, Thomas J. Cunningham

**Author notes:** These authors contributed equally to this work (David Thompson, Chloe Williams). Senior authors (Elizabeth M.C. Fisher, Thomas J. Cunningham).

## Abstract

Amyotrophic lateral sclerosis (ALS) caused by mutation in *superoxide dismutase 1* (*SOD1*) accounts for 15-30% of familial ALS and is typically autosomal dominant. How single base pair/amino acid changes in this small protein cause neurodegeneration is unknown. In North America, SOD1^A4V^ is the most common familial ALS SOD1 mutation and results in an aggressive form of ALS. Here, we present a novel genomically humanised mouse model of *SOD1^A4V^*, in which the mouse *Sod1* locus has been replaced by the human *SOD1* gene, with intact genomic architecture of exons and introns, but bearing an A4V mutation. In agreement with previously reported human genomic knock-in mice, the phenotype is mild; however, transcriptomic and metabolomic profiling reveal significant dysregulation of glycolysis, the tricarboxylic acid (TCA) cycle, and lipid metabolism. These changes suggest an early bioenergetic imbalance that precedes neuromuscular impairment. Our findings support metabolic dysfunction as an early event in ALS pathogenesis. This freely available *SOD1^A4V^* model provides a valuable tool for studying ALS progression and identifying therapeutic targets for pre-symptomatic treatment.

**SUMMARY STATEMENT:** This study describes the generation and analysis of novel genomically humanised *SOD1^A4V^* mice, revealing metabolic dysfunction through integrated multi-omic analyses, characterising a freely available potential pre-symptomatic ALS model for future research.

## INTRODUCTION

Amyotrophic lateral sclerosis (ALS) is a fatal neurodegenerative disease characterised by the progressive loss of both upper and lower motor neurons, leading to muscle weakness, wasting and progressive paralysis. It is the most common cause of motor neuron disease globally, with an incidence of approximately 2/100,000 population and up to a 1/300 lifetime risk of disease (Al-Chalabi & Hardiman, 2013; Chiò et al., 2013). The aetiology of ALS is not understood. The age of onset is around 60 years old for sporadic ALS (sALS), which arises without a known family history, and 40-60 years old for hereditary cases of familial ALS (fALS), with a median survival of 3-5 years from onset (Ingre *et al*., 2015). There are currently no treatments that halt the progression of the disease and a paucity of biomarkers for disease predisposition and pre-symptomatic onset.

ALS likely develops through an interplay of genetic susceptibility and largely unknown environmental factors. Approximately 90% of cases are sALS; the remaining ∼10% are fALS, of which 70% have a mutation identified in an ‘ALS gene’, most of which are autosomal dominantly inherited (Saeed et al., 2009). Over 50 different ALS associated genes are known, with mutations in *SOD1, FUS, TARDBP* and *C9ORF72* causal for 55% of European fALS cases (Mejzini *et al*., 2019; Zou *et al*., 2017; Renton *et al*., 2014). Thus, ALS is a clinically defined set of diseases with different origins that result in the death of upper and lower motor neurons (Magnussen & Glass, 2017).

Mutations in *superoxide dismutase 1* (*SOD1*) were the first genetic cause of ALS to be discovered and account for 15-30% of fALS and 2% of sALS cases (Zou *et al*., 2017; Saccon et al., 2013; Rosen *et al*., 1993; Deng et al., 1993). Disease-causing mutations have been identified in each of the 5 exons of this gene and over 180 different potentially pathogenic variants have been recognised (Abel *et al*., 2013). Genotype-phenotype associations show patients carrying A4V, G85R and G93A mutations typically have rapid disease progression. Other variants, such as P66S and G16S, are associated with an earlier age of onset (Mejzini *et al*., 2019). In part due to a founder effect, the A4V variant located in exon 1 makes up ∼50% of SOD1 pathogenic variants in North America and has an average survival time of less than 2 years (Saeed *et al*., 2009). Note that using contemporary nomenclature, the A4V mutation is the in the fifth codon position (A5V), but previously the methionine start codon was not counted – for historical consistency we refer to this mutation as A4V.

The relative frequency of the A4V variant accounts for its representation within pre-symptomatic familial ALS research programs especially in North America. Some programs are undertaking detailed review of genetically predisposed individuals to assess features that may mark the shifts from pre-symptomatic to disease onset, i.e. the transition from a clinically silent period where biomarkers of disease are present to a prodromal phase of non-impactful clinical features (Phenotransition) (Benatar *et al*., 2024). The conversion of prodromal clinical features into the symptomatic expression of classifiable ALS and the onset of the clinically manifest stages may also be tracked (Phenoconversion) (Benatar *et al*., 2022, Benatar *et al*., 2023). However, robust markers of phenotransition and phenoconversion remain to be found and *SOD1*-ALS disease aetiology is unclear.

Genetically modified mouse models have been of great utility for ALS research since the generation of transgenic *SOD1^G93A^*(*tg.SOD1^G93A^*) mice that overexpress mutant human *SOD1* in a background of endogenous mouse *Sod1* expression (Gurney *et al*., 1994). Hemizygous *tg.SOD1^G93A^* mice carry ∼twenty-five transgene copies, and exhibit ∼4-fold higher total SOD1 protein levels in the brain versus non-transgenic mice, and display early onset (∼90 days of age) and fast progression of disease (∼60 days). However, presymptomatic or early-stages of pathogenic processes are difficult to assess in such rapidly progressing models, and spurious phenotypes can arise from overexpression *per se*, rather than the effect of the introduced mutation (De Giorgio *et al*., 2019; Saccon et al., 2013). Transgenic *SOD1^A4V^*(*tg.SOD1^A4V^*) mice, generated in the same study but not widely used, have substantially lower total SOD1 protein levels (only 50% higher than non-transgenic mice) and show no overt phenotype suggesting a threshold level of SOD1 may be required for motor phenotypes to develop in mouse (Gurney et al., 1994; Deng *et al*. 2006).

Subsequent SOD1 mouse models have aimed to recapitulate human ALS by using knock-in of pathogenic variants within the endogenous mouse *Sod1* gene to mimic physiological levels of SOD1 expression (De Giorgio *et al*., 2019). Without overexpression, typically the onset and progression of disease has been less aggressive, despite using variants that accentuate these features in human ALS (De Giorgio *et al*., 2019; Fisher & Bannerman, 2019).

Recently, we developed a genomically humanised *hSOD1^WT/WT^* gene targeted mouse strain in which the mouse *Sod1* gene was replaced with the wildtype human *SOD1* gene at the endogenous *Sod1* locus (Devoy *et al*., 2021). The human *SOD1* genomic region extends from the ATG start codon to the end of the 3′-UTR; this freely available *hSOD1^WT/WT^* strain retains upstream regulatory sequences including the promotor of mouse *Sod1*, allowing the study of wild-type human SOD1 at physiological expression levels. The *hSOD1^WT/WT^* mouse strain shows no adverse phenotypes over a normal lifespan compared to non-transgenic littermates. Here, we take our genome engineering a step further and describe the generation of a new genomically humanised *SOD1* knock-in mouse, carrying the *hSOD1^A4V^*variant. This new strain affords the opportunity to model heterozygous and homozygous effects of a clinically important mutation.

Heterozygous *hSOD1^WT/A4V^* and homozygous *hSOD1^A4V/A4V^* mice have reduced SOD1 protein levels and SOD1 enzyme activity. Over the lifespan of these mice, up to 700 days, *hSOD1^A4V^* animals are not distinguishable from wild type mice in neuromuscular assessments. However, longitudinal transcriptomic and metabolic profiling identified dysregulation of key pathways in energy metabolism, with glycolysis, the TCA cycle, and lipid utilisation being perturbed in both *hSOD1^WT/A4V^* and *hSOD1^A4V/A4V^* animals. This novel humanised model will be useful for examining the factors that promote the transition from an asymptomatic, genetically predisposed state to clinical disease, allowing insight into the clinically silent pre-manifest stage of asymptomatic ALS (Benatar et al., 2023; Benatar et al., 2024). This mouse line is freely available from the European Mutant Mouse Archive (EMMA; to be added).

## RESULTS

### Generation and initial characterisation of genomically humanised *SOD1^A4V^* mice

Using CRISPR/Cas9 mutagenesis, we edited our previously published *hSOD1^WT/WT^* mouse strain (Devoy *et al*., 2021) to generate SOD1-A4V (*hSOD1^A4V^*) mice (Fig. 1A). Note that the A4V mutation was placed into our existing *hSOD1^WT^* allele, in which we had created a duplication of the exon 4-5 region, where several important SOD1 variants occur; in the future, this will allow the effects of mutations in these two exons to be temporally and spatially analysed, using Cre recombinase.

**Figure 1.**
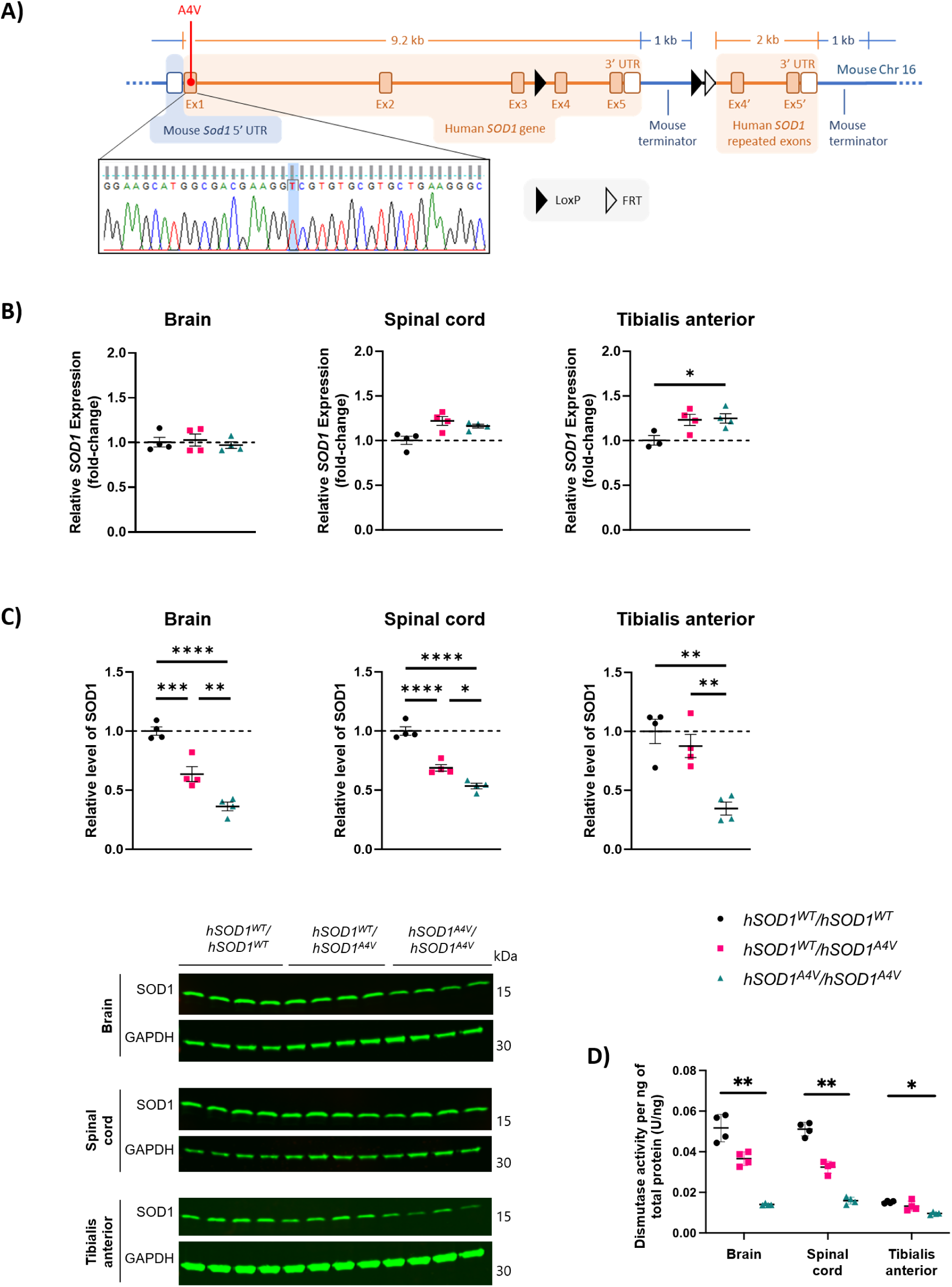
SOD1 expression and activity in *hSOD1^A4V^* mice. A) Schematic of the *hSOD1^A4V^* allele. Humanised sequence is indicated in orange, mouse sequence in blue. Ex, exon; UTR, untranslated region. Inset: Sequencing trace from a homozygous *hSOD1^A4V^* animal showing A4V point mutation (C->T; highlighted). B) Dot plot showing relative expression of total *SOD1* mRNA using conserved mouse-human *SOD1/Sod1* primers in tissues from 3-month-old male animals (n=3-4 animals per genotype per tissue). All values are expressed as a fold change relative to the average *SOD1* expression in *hSOD1^WT/WT^* control animals. Data normalised to *GAPDH.* C) Western blots using an anti-SOD1 antibody (aa131-153, reacts with mouse and human) in tissue extracts from 3-month-old male animals. Human SOD1 migrates with a higher apparent molecular mass than mouse SOD1 in SDS-PAGE. Anti-SOD1 signal intensity was normalized to anti-GAPDH signal. Intensity in the same lane. All values are expressed as a fold change relative to the average SOD1 level in *hSOD1^WT/WT^* control animals. n=4 per genotype per tissue. D) Dot plot showing spectrophotometric dismutase activity assays in tissues from male 3-month-old *hSOD1^A4V^* mice. One-way ANOVA with Tukey’s post hoc test. \**P* < 0.05, \*\**P* < 0.01, \*\*\**P* < 0.001, \*\*\*\**P* < 0.0001.

However, as the A4V mutation is in exon 1, we did not use this feature here.

We analysed animals homozygous for the wild-type humanised locus (*hSOD1^WT/WT^*) as controls, compound heterozygotes carrying the wild-type humanised locus and the humanised A4V allele (*hSOD1^WT/A4V^*), or homozygous for the humanised A4V allele (*hSOD1^A4V/A4V)^* (herein, the use of *hSOD1^A4V^* denotes heterozygote and homozygote A4V animals). To determine the effect of A4V mutation on *SOD1* mRNA expression, we performed RT-qPCR in a range of tissues (brain, spinal cord (SC), quadriceps, tibialis anterior (TA), extensor digitorum longus (EDL), soleus, kidney, and liver) from 3-month-old male and female *hSOD1^A4V^* animals and *hSOD1^WT/WT^*littermate controls. *SOD1* mRNA expression did not differ significantly between *hSOD1^A4V^* animals and *hSOD1^WT/WT^* littermate controls in the majority of tissues examined (Fig. 1B and S1) although *SOD1* mRNA expression was increased in the TA muscle of male *hSOD1^A4V/A4V^* mice (Fig. 1B) and the liver of male *hSOD1^WT/A4V^*mice (Fig. S1A), compared to *hSOD1^WT/WT^*.

In contrast to mRNA levels, *hSOD1^A4V^* animals had significantly reduced SOD1 protein levels in almost all tissues examined in both sexes (Fig. 1C, S2 and S3). SOD1 protein levels correlated with the zygosity of the A4V mutation, with reduced SOD1 detected in *hSOD1^A4V/A4V^* versus *hSOD1^WT/A4V^* animals. Protein expression in *hSOD1^A4V/A4V^* animals ranged from half to approximately one tenth of the levels found in tissues from *hSOD1^WT/WT^*littermate controls with *hSOD1^WT/A4V^*showing intermediate levels SOD1 activity in homozygous *hSOD1^A4V/A4V^* tissues was also significantly reduced in all tissues examined from 3-month-old male and female animals, exhibiting between 10-40% of the activity observed in the corresponding tissue from *hSOD1^WT/WT^*littermates (Fig. 1D and S4).

To determine the effect of the A4V mutation on SOD1 stability, we assessed levels of soluble disordered SOD1 in tissues from 12-month-old *hSOD1^A4V^* and *hSOD1^WT/WT^* littermates (Fig. 2A-D). The highest levels were detected in brain and were significantly increased in brain from *hSOD1^WT/A4V^* and in brain and liver from *hSOD1^A4V/A4V^* animals (Fig. 2A, 2D).

**Figure 2.**
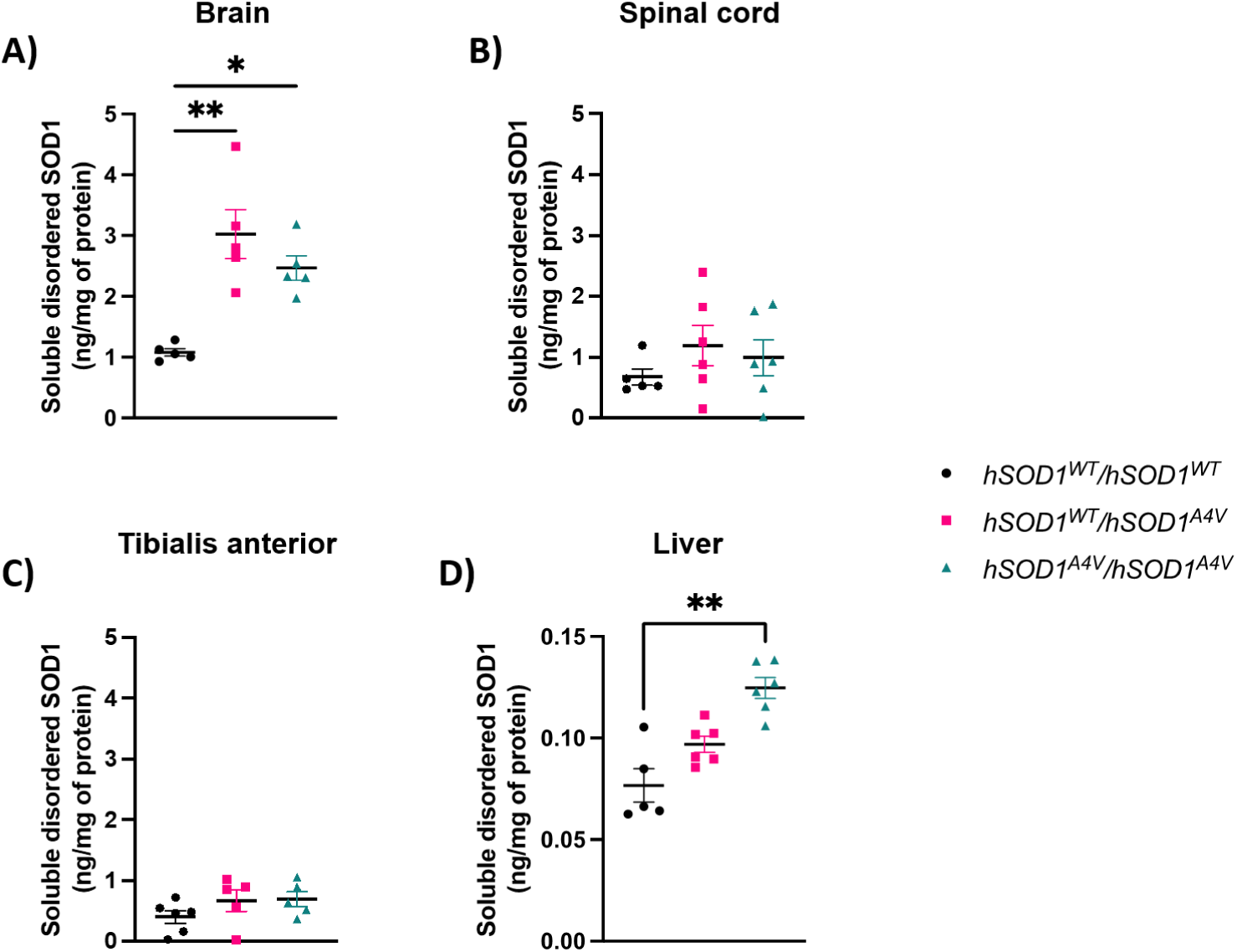
Disordered SOD1 in *hSOD1^A4V^* mice. Levels of soluble disordered SOD1 in supernatants of A) brain, B) spinal cord, C) tibialis anterior and D) spinal cord of 12-month-old male and female animals, quantified by ELISA performed with a peptide antibody against amino acids 24-39 recognizing an epitope exposed in disordered SOD1. One-way ANOVA with Tukey’s post hoc test. \**P* < 0.05, \*\**P* < 0.01.

### *hSOD1^A4V^* mice lack an overt neurodegeneration phenotype

We performed extensive motor and behavioural phenotyping of *hSOD1^A4V^* animals. Onset of progressive motor dysfunction is the hallmark of human ALS and is shown in overexpression transgenic ALS mouse models by deficits in grip strength, Rotarod and gait (Jaarsma et al., 2000; Filali et al., 2011; Mancuso et al., 2011; Hadzipasic et al., 2014; Oliván et al., 2015). However, we did not observe deficits in these tests in our knock-in *hSOD1^A4V^* mice (Fig. S5), nor did we detect atrophy of the TA muscle (Fig. S6). We also did not detect NMJ denervation in the TA, soleus or lumbrical muscles of *hSOD1^A4V^* animals (Fig. S7A-D), nor loss of motor neurons in the lumbar L3-L5 region of the SC, suggesting that the *SOD1-A4V* mutation does not impact motor neuron survival in *hSOD1^A4V^* mice at the ages studied (Fig. S7E). Furthermore, in contrast to *tg.SOD1^G93A^* mice (Tosolini et al., 2022), we did not observe *in vivo* axonal transport deficits in TA-innervating motor axons (Fig S8).

### *hSOD1^A4V^* mice have mild metabolic phenotypes

Metabolic changes have been widely observed in ALS mouse models, including at pre-symptomatic stages (Bame et al., 2014; Henriques et al., 2015; Fernández-Beltrán et al., 2021; Lee et al., 2021; Maksimovic et al., 2023); hence, we sought to establish if metabolism is perturbed in *hSOD1^A4V^* mice. To assess the impact of the A4V mutation on bodyweight, *hSOD1^A4V^* animals were weighed monthly from 1 to 26 months of age (Fig. 3A, 3B). There was a significant effect of age (p = <0.001) and genotype (p = 0.022), and a significant interaction between age and genotype (p = 0.003) on bodyweight in female *hSOD1^A4V^* animals (Fig. 3A). Female *hSOD1^A4V/A4V^*mice gained weight more slowly than controls after 6 months of age, suggesting this difference was not due to impaired development. No difference in bodyweight was observed in male animals (Fig. 3B).

**Figure 3.**
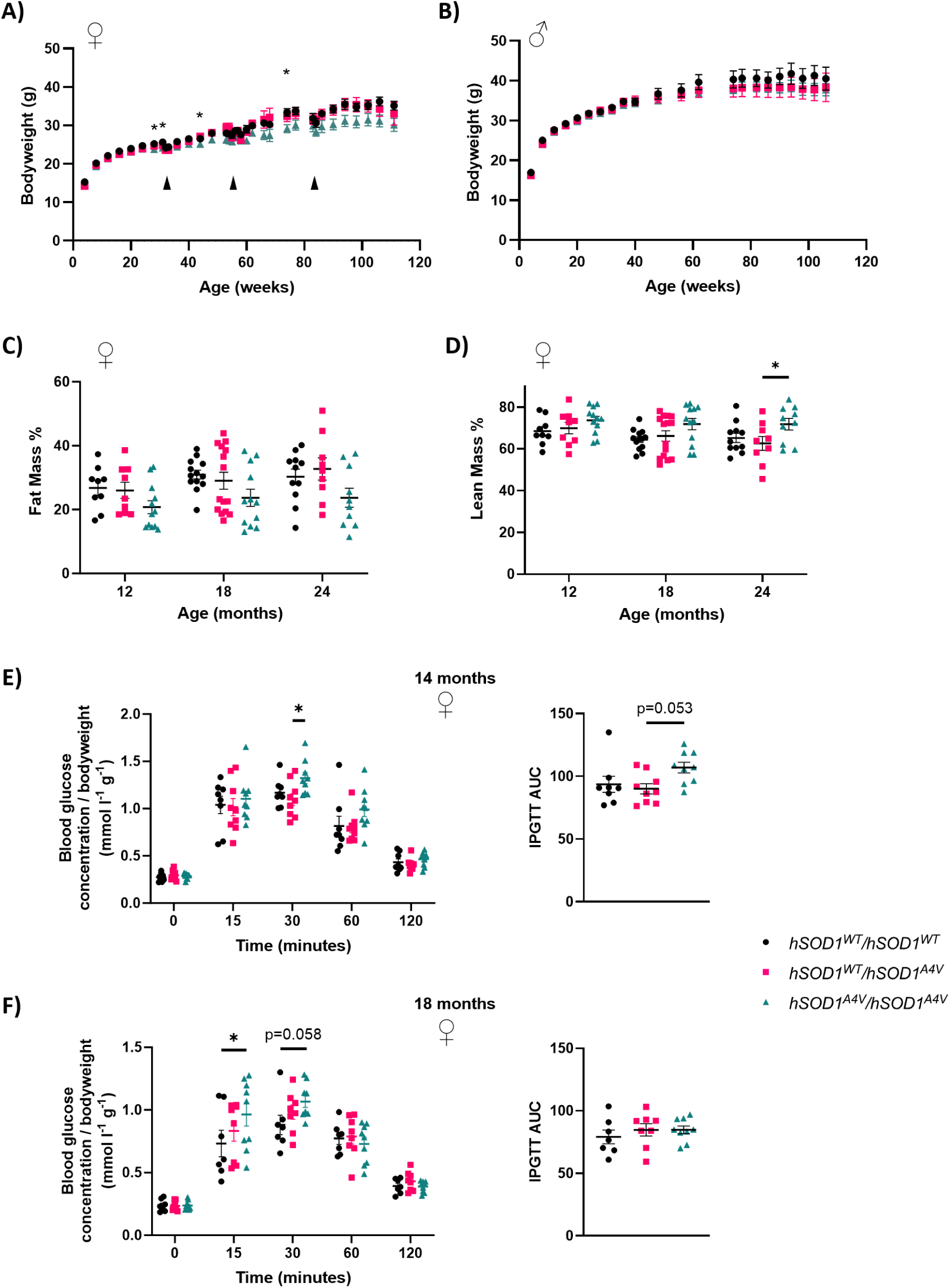
*hSOD1^A4V^* mice have a mild metabolic phenotype. (A, B) Monthly bodyweight measurement data from male (n=12-16 per genotype at 4 weeks old, n=8-12 per genotype at 106 weeks old) and female (n=14-16 per genotype at 4 weeks old, n=6-9 per genotype at 111 weeks old) *hSOD1^A4V^* mice. Arrowheads indicate ages where automated wheel running experiments were performed, showing expected dips in bodyweight. Mixed effects analysis followed by the Tukey multiple comparisons test. (C, D) Echo MRI data showing fat mass and lean mass as a percentage of total bodyweight in female (n=9-15 per genotype) *hSOD1^A4V^* animals. Two-way ANOVA with Tukey’s post hoc test. (E, F) Intraperitoneal glucose tolerance testing (IPGTT) data from (E) 14-month-old (n=8-9 per genotype) and (F) 18-month-old (n=7-9 per genotype) female *hSOD1^A4V^* mice. Changes in blood glucose over time were analysed using two-way repeated measures ANOVA followed by Tukey’s test. Area under the curve (AUC) for each graph was analysed using one-way ANOVA followed by Tukey’s test. Reported significance values in all panels are from Tukey’s test. Main effects are reported in the main text. Mean ± SEM displayed in all panels. \**P* < 0.05.

To examine whether reduced bodyweight in female *hSOD1^A4V^*animals could be attributed to changes in fat or lean mass we performed ECHO-MRI on 12-18-, and 24-month-old animals (Fig. 3C and D). 2-way ANOVA revealed a significant effect of genotype, but not age, on percentage fat mass (p = 0.002) and percentage lean mass (p = 0.002) in female *hSOD1^A4V^* animals. Pairwise comparisons conducted at each age found a significant increase in percentage lean mass in 24-month-old *hSOD1^A4V/A4V^* compared to *hSOD1^WT/A4V^*females. However, across all ages there was only a general trend towards decreased percentage fat mass and increased percentage lean mass in female homozygous *hSOD1^A4V/A4V^* mice compared to the other groups (age and sex matched).

Glucose metabolism was investigated in 14-and 18-month-old female *hSOD1^A4V/A4V^* mice using the intraperitoneal glucose tolerance test (IPGTT; Fig. 3E and 3F). Multiple pairwise comparisons were conducted at each sampling time. A significant increase in blood glucose was detected 15 minutes and 30 minutes after glucose infusion in 14-and 18-month-old female *hSOD1^A4V/A4V^*animals, respectively, compared to *hSOD1^WT/WT^* littermate controls, suggesting an impaired response to glucose.

### Transcriptomic changes at different timepoints in *hSOD1^A4V^* mice

We performed QuantSeq 3′ mRNA sequencing (Lexogen) in the TA muscle and lumbar SC of 6-, 12-and 20-month-old male animals and assessed gene expression differences in *hSOD1^WT/A4V^* and *hSOD1^A4V/A4V^* animals versus *hSOD1^WT/WT^* littermate controls. Average gene expression was obtained (n=4 animals in each genotype/time point) and the proportion of differentially expressed genes (DEGs) between groups was initially determined using a false discovery rate adjusted p-value of < 0.1 as the cut-off. A small number of significant DEGs were identified (Table S1) in the TA muscle of *hSOD1^WT/A4V^* and *hSOD1^A4V/A4V^*animals at 6 and 20 months of age and in the lumbar SC of *hSOD1^WT/A4V^*and *hSOD1^A4V/A4V^* at 20 months of age, compared to *hSOD1^WT/WT^* littermate controls (Table S1, Fig. S9 and S10). To widen the search in order to identify biological pathways, analysis was then conducted using a P-value of <0.01 (Fig. S9 and S10). There was minimal overlap of DEGs between genotypes and time points in within-tissue comparisons in the TA and SC (Fig. S11, Table S2).

DEGs with the largest fold change (10 most upregulated and 10 most downregulated genes) were identified in each data set (Table S3, for a complete list of DEGs see Tables S4 and S5). A large proportion of these genes were predicted or unannotated (26% of genes for TA, 50% of genes for SC). Of the annotated genes, several have been previously linked to ALS (Table S3).

To characterise the functional relevance of the observed transcriptional changes, the enrichment analysis tool Enrichr (Chen et al., 2013; Kuleshov et al., 2016; Xie et al., 2021) was used to identify gene ontology (GO) terms and infer canonical pathways regulated by DEGs.

GO enrichment analysis across DEGs at 6-, 12-and 20-months in the TA of *hSOD1^WT/A4V^* and *hSOD1^A4V/A4V^* animals, compared to *hSOD1^WT/WT^* littermate controls, revealed altered expression of genes involved in: aerobic electron transport chain activity; mitochondrial function; malate and nitric oxide metabolism; ribosomal function; stress granule assembly; among others (Fig. S12, S13). GO enrichment analysis across DEGs at 6-, 12-and 20-months in the SC of *hSOD1^WT/A4V^*and *hSOD1^A4V/A4V^* animals, compared to *hSOD1^WT/WT^*littermate controls, revealed GO terms related to: oxidative stress-induced cell death; ATPase complex activity; oxidoreductase activity; malate and pyruvate metabolism; among others (Fig. S14, S15). DEGs involved in aerobic respiration and translation were detected in the TA and SC of *hSOD1^WT/A4V^*and *hSOD1^A4V/A4V^* animals, suggesting that despite few overlaps between DEGs, the pathways that are affected are common upon *SOD1^A4V^*mutation.

The presence of commonly altered pathways is further shown in the enriched KEGG pathways. Of the 10 most significant identified KEGG pathways, 7 are represented in both the TA and SC of *hSOD1^WT/A4V^* when compared to *hSOD1^WT/WT^* littermate controls (Fig. 4A, 4B). These pathways include oxidative phosphorylation, ribosome, electron transport chain, and *Wnt* signalling. Several KEGG pathways related to neurodegenerative disease were also identified in the *hSOD1^WT/A4V^*TA and SC, including the terms: pathways of neurodegeneration; Alzheimer’s disease; Parkinson’s disease; prion disease. Additional altered KEGG pathways included: energy metabolism including glycogen metabolism and glycolysis/gluconeogenesis, which were identified in the TA and SC, respectively.

**Figure 4.**
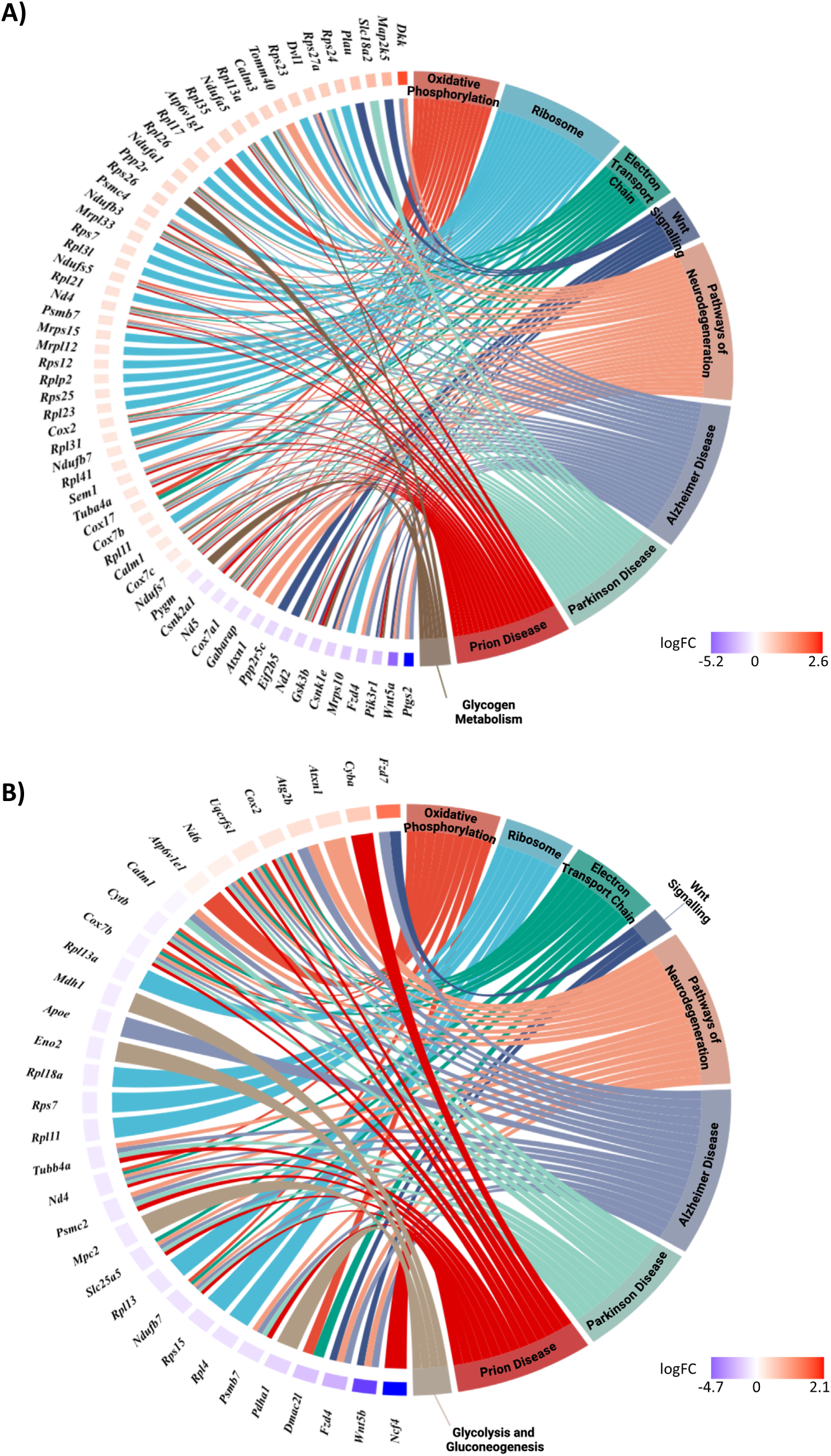
Chord diagram showing the core genes of significant affected pathways in *hSOD1^WT/A4V^*mice compared to littermate controls. Significant pathways identified from DEGs in the A) TA and B) SC of *hSOD1^WT/A4V^* vs *hSOD1^WT/WT^* 6-, 12-and 20-month-old male mice. Significant pathways are shown on the right and the fold change of core genes is shown on the left. Left-right connections indicate gene membership in a pathways leading-edge subset. Image constructed using Biorender.

Enrichment analysis identified few significantly altered KEGG pathways in *hSOD1^A4V/A4V^* tissues; however, pyruvate metabolism was enriched in both the TA and SC of *hSOD1^A4V/A4V^* mice (Fig. S16).

Transcriptomic analysis of changes in the TA and SC of *hSOD1^WT/A4V^* and *hSOD1^A4V/A4V^*animals compared to *hSOD1^WT^*^/*WT*^ littermate controls highlighted several pathways related to neurodegenerative disease and significant changes in metabolism, encompassing genes related to glucose and pyruvate metabolism, the electron transport chain and ATPase complex activity.

### Metabolic changes over life in *hSOD1^A4V^* mice

To investigate potential metabolomic changes indicated from metabolic phenotyping and transcriptomic analyses, we performed LC-MS and GC-MS on plasma from 4-, 6-, 9-, 13-and 20-month-old *hSOD1^A4V^* male and female mice (Fig. S17-19) and tissue samples (SC, TA and liver) from 20-month-old *hSOD1^A4V^* male mice (Fig. S20 and S21).

Univariate analysis identified a total of 3, 15, 1 and 8 significantly altered metabolites in plasma collected at 4-, 6-, 9-and 20-months of age, respectively, in *hSOD1^WT/A4V^* mice compared to *hSOD1^WT/WT^* littermate controls (Fig. S17-19). *hSOD1^WT/A4V^* animals had increased levels of hydroxy fatty acids, lysophospholipids, acylcarnitines, and amino-and keto-acids, and reduced levels of glucose and the NAD precursor niacinamide. Interestingly, threonic acid (also known as dehydroascorbic acid) was significantly decreased in the blood of 6-month-old *hSOD1^WT/A4V^* mice compared to *hSOD1^WT/WT^* littermate controls. Threonic acid is produced upon oxidation of ascorbic acid, a key antioxidant with an established protective role against oxidative stress in the CNS (Rice, 2000).

Univariate analysis identified a total of 2, 6, 3 and 9 significantly altered metabolites in plasma collected at 6-, 9-, 13-and 20-months of age, respectively, in *hSOD1^A4V/A4V^* mice compared to *hSOD1^WT/WT^* littermate controls (Fig. S17-19). *hSOD1^A4V/A4V^* animals had increased levels of amino acids and their derivatives, increased lysophospholipids, and reduced levels of fatty acids and sterols. Pyruvic acid, a central metabolite in glycolysis, was increased. The beta-hydroxybutyrate ester (3-hydroxybutyric acid), currently under trial for treatment of ALS patients (Ludolph, 2024), was significantly decreased.

Univariate analysis identified a total of 0, 4 and 4 significantly altered metabolites in SC, TA and liver, respectively, in 20-month-old *hSOD1^WT/A4V^* mice compared to *hSOD1^WT/WT^* littermate controls (Fig. S20 and S21). In TA, the nucleoside guanosine and in the dinucleotide NAD were decreased (Fig. S20B). In liver, the TCA cycle intermediate succinic acid was increased, as was the pentose phosphate pathway intermediate ribose-5-phosphate (Fig. S21).

Univariate analysis identified a total of 2, 2, and 9 significantly altered metabolites in SC, TA and liver, respectively, in *hSOD1^A4V/A4V^*mice compared to *hSOD1^WT/WT^* littermate controls (Fig. S20 and S21). In SC, the lysophospholipid lysoPE(0:0/18:3(6Z,9Z,11Z)) was decreased whereas hippuric acid was increased (Fig. S20A). Sorbose, a keto-hexose sugar necessary for vitamin C biosynthesis, was significantly increased (Fig. S20B). In liver, several amino acid and glycine conjugates were significantly increased, and ribose-5-phosphate was significantly increased, in *hSOD1^A4V/A4V^* mice compared to *hSOD1^WT/WT^* littermate controls (Fig. S21).

Reduced glutathione (GSH) reacts with reactive oxygen species and is converted to its oxidised form (GSSG) in the process – forming a key part of the antioxidant defence (Owen & Butterfield, 2010). A disruption in SOD1 function has been shown to perturb other antioxidant systems, hence we examined the levels of GSH and GSSG in blood from 4-and 20-month-old *hSOD1^A4V^* mice (Park et al., 2023; Wang et al., 2011). Levels of GSH were unchanged at both timepoints (Fig. S22).

However, we found a significant decrease in GSSG in 13-and 20-month-old *hSOD1^A4V/A4V^* animals compared to controls. Accordingly, we observed an increase in the ratio of GSH-to-GSSG in 20-month-old *hSOD1^A4V/A4V^* animals.

### Integrated transcriptomic and metabolomic analysis identifies changes in energy metabolism

Integrated analysis of significant DEGs (across 6-, 12-and 20-month timepoints) and significantly changed metabolites (in blood collected at 4-, 6-, 9-, 13-and 20-months and the TA and SC at 20-months) highlighted energy metabolism pathways encompassing glycolysis, lipid utilisation and the TCA cycle in both *hSOD1^WT/A4V^* and *hSOD1^A4V/A4V^* mouse models (Fig. 5 and S23).

**Figure 5.**
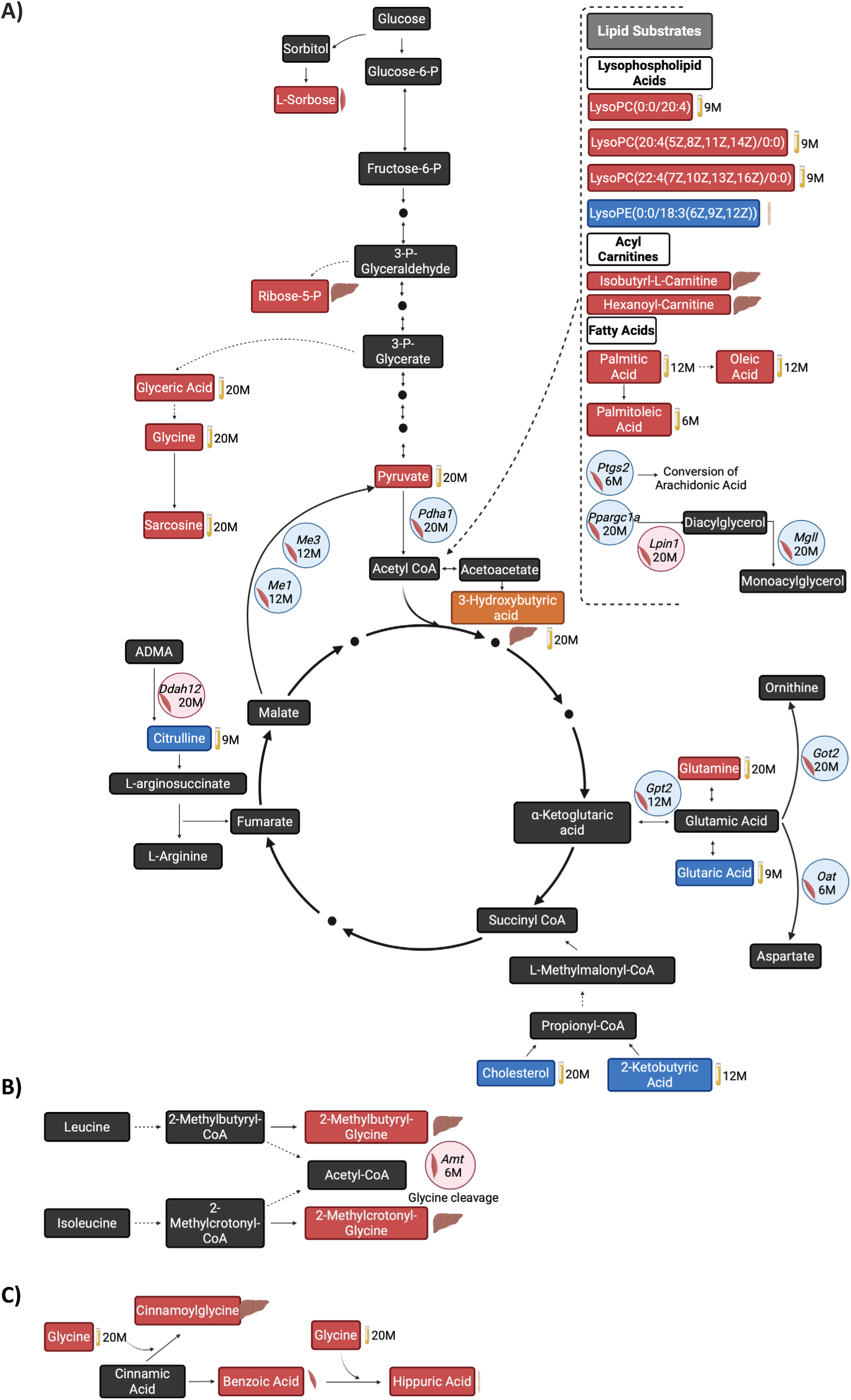
Integrated transcriptomic and metabolomic analysis of *hSOD1^A4V/A4V^* mice. Schematic representation of dysregulated pathways. A) energy metabolism pathways, B) leucine and isoleucine degradation, C) aromatic acid detoxification in *hSOD1^A4V/A4V^* mice. Metabolites are depicted by rectangles, red: increased, blue: decreased and black: undetected. Genes are depicted by circles, red: increased, blue: decreased. Solid arrows indicate direct reaction, dotted arrows indicate the reaction is broken by one or more sub-reactions. The tissue in which genes or metabolites were significantly altered are depicted by the presence of a muscle, spinal cord or liver image. The timepoint at which a gene or metabolite was detected as significantly changed is indicated in months (M). Image constructed using Biorender.

Integrated analysis of significantly changed genes and metabolites in *hSOD1^WT/A4V^* animals produced a model in which the majority of identified genes encode key enzymatic intermediates of glycolysis and the citric acid cycle. Genes encoding enzymes involved in pyruvate production and transport were downregulated as were genes encoding enzymes involved in the production of fructose-6-phosphate. Metabolites highlighted through integrated analysis included increased levels of glucose together with increased levels of lipid substrates (Fig. S23). Despite downregulation of genes related to glycolysis, there is an enrichment of upregulated genes involved in the formation and activity of complexes I, IV and V of the electron transport chain (Fig. S24A). Integrated analysis of significantly changed genes and metabolites in *hSOD1^WT/A4V^*compared to *hSOD1^WT/WT^* littermate controls identified dysregulation in additional pathways including xanthine and NAD metabolism (Fig. S24B-C).

Integrated analysis of significant DEGs and metabolites in *hSOD1^A4V/A4V^*animals produced a model highlighting several metabolic pathways including lipid utilisation and auxiliary TCA metabolism pathways (Fig. 5). The model shows an increase in the central glycolysis metabolite pyruvate in *hSOD1^A4V/A4V^* animals compared to *hSOD1^WT/WT^* littermate controls, whilst genes encoding enzymes for the conversion of malate into pyruvate and pyruvate into acetyl-co-A are downregulated. Lipid substrates for metabolism, including acyl-carnitines and fatty acids, are increased. The asymmetric dimethylarginine (ADMA) degradation pathway, linked to the urea cycle and fumarate production, is implicated in the model. The gene encoding the enzyme responsible for converting ADMA to citrulline, *Ddah12*, is significantly upregulated whilst citrulline is significantly decreased. Sarcosine metabolism is also highlighted, with increased levels of glyceric acid, glycine and sarcosine detected in the blood of 20-month animals compared to *hSOD1^WT/WT^* littermate controls. The glutamate-glutamine cycle also shows up with significant DEGs and metabolites in *hSOD1^A4V/A4V^*animals. The gene, *Gpt2*, encoding an enzyme involved in the conversion of α-ketoglutaric acid to glutamic acid is downregulated as are genes encoding enzymes involved in the conversion of glutamic acid to ornithine and aspartate. Further, glutamine is significantly increased whilst glutaric acid is significantly decreased (Fig. 5A). Integrated analysis of significant DEGs and metabolites in *hSOD1^A4V/A4V^*compared to *hSOD1^WT/WT^* littermate controls highlighted dysregulation in additional pathways including the leucine and isoleucine degradation pathway and aromatic acid detoxification (Fig. 5B and 5C).

The integrated analyses of significant DEGs and metabolites in *hSOD1^A4V^*animals compared to *hSOD1^WT/WT^* littermate controls suggests *SOD1^A4V^* mutation results in dysregulation of glycolysis, the TCA cycle and lipid utilisation in both the *hSOD1^A4V^* heterozygous and homozygous mouse models. Further, the electron transport chain is also highlighted by the presence of dysregulated gene expression in *hSOD1^WT/A4V^* animals whilst auxiliary TCA pathways are defined in the *hSOD1^A4V/A4V^* animals. Overall, the integrated analysis of our transcriptomic and metabolomic data suggests a metabolic signature of disease in the absence of overt neurodegeneration.

## DISCUSSION

Here, we present the characterization of the first genomically humanized SOD1 knock-in mouse model of ALS, offering insights into potential metabolic perturbations associated with SOD1-ALS mutation in the absence of neurodegeneration. Using our knock in humanised model we were able to study heterozygous and homozygous mutation effects. This is important as in addition to SOD1 gain of function (GOF) phenotypes, SOD1-ALS patients may have some degree of SOD1 loss of function (LOF), because enzymatic activity is typically reduced in the majority of variants (Saccon et al., 2013). Furthermore, SOD1 LOF has a more severe impact on neurological function in humans than in mice (Andersen et al., 2019; Park et al., 2019; Park et al., 2023; Saccon et al., 2013), potentially because *SOD1* is an essential gene in humans whereas this is not the case in mice (Blomen et al., 2016; Dickinson et al., 2016). Therefore, both GOF and LOF phenotypes remain relevant for ALS disease modelling.

We observed reduced SOD1 levels and activity in *hSOD1^A4V^*mice, which is consistent with results from SOD1-A4V patients (Kiskinis et al., 2014; Bowling et al., 1993; Saccon et al., 2013). We found particularly low SOD1 activity in skeletal muscle of *hSOD1^A4V^* mice, especially when compared to wildtype *mSod1*/*mSod1* mice (i.e. both mouse *Sod1* genes intact) (Fig. S25). This may be due to the compounded effects of lower SOD1 transcription in muscle in *hSOD1^WT/WT^*mice versus *mSod1*/*mSod1* (Devoy *et al*., 2021), plus the decreased stability of SOD1-A4V (vs SOD1-WT) in *hSOD1^A4V^* mice. The minimum level of SOD1 required to prevent LOF phenotypes in mice is unknown, and it is possible that LOF phenotypes may be observed in *hSOD1^A4V^*mice, and potentially SOD1-A4V patients.

The presence of aggregated SOD1 protein species, which arise from the misfolding and accumulation of soluble disordered SOD1, is a hallmark of ALS observed in SOD1-ALS patients and mouse models (Shibata et al., 1996; Shibata et al., 1996; Kato et al., 2000; Bruijn et al., 1998).

Increased levels of soluble disordered SOD1 were detected in the tissues of *hSOD1^A4V^* animals, in the absence of protein aggregates, potentially suggesting a pre-manifest prodromal state of disease (Benatar et al., 2023; Benatar et al., 2024). In mice, high levels of SOD1 protein are found in the brain and liver, providing an explanation for a significant increase in disordered SOD1 detected in these tissues. We did not observe development of motor symptoms in *hSOD1^A4V^*mice, suggesting that expression of the A4V mutation at physiological levels may be insufficient to drive the classical ALS phenotype observed in transgenic overexpression models, within the lifespan of this strain of mouse. These findings are consistent with those of the low copy transgenic *tg.SOD1^A4V^* strain (Gurney et al., 1994), where a low level of mutant SOD1 protein and lack of protein aggregation were accompanied by an absence of a motor phenotype. However, a phenotype was observed when *tg.SOD1^A4V^* mice were crossed with a line overexpressing wild-type hSOD1 at a high levels, suggesting that wild type SOD1 can act as a substrate for aggregation and that total hSOD1 levels are a limiting factor for neurodegeneration in models expressing mutant SOD1 transgenes (Deng et al., 2006).

To our knowledge, there is only one other SOD1 A4V mutant mouse, a *Sod1^A4V^* knock-in strain in which the A4V mutation has been introduced into the mouse *Sod1* gene (JAX 031341). This strain also has no documented phenotype on the JAX strain description. A small number of different low-copy transgenic SOD1-ALS models exist that show motor phenotypes late in life (Deng et al., 2006; Gurney et al., 1994 Deitch et al., 2014; Watanabe et al., 2005), but there has been no direct comparison of protein levels between these different models, noting that transgenic copy number is not directly proportional to translational output (Acevedo-Arozena et al., 2011). Thus, the simplest explanation for a lack of motor phenotypes in A4V models is low SOD1 levels and thus insufficient disordered SOD1 to initiate fulminant aggregation and the likely related neurotoxic cascade.

Several observations support a mild metabolic phenotype in *hSOD1^A4V^*mice, which could be linked to gain or loss of SOD1 function, or both. There are similarities between the phenotypes observed in *hSOD1^A4V^*and published observations from *Sod1*^-/-^ mice, which may indicate a SOD1 LOF effect. Female *hSOD1^A4V/A4V^* animals had reduced bodyweight versus *hSOD1^WT/WT^* littermate controls and a trending decrease in fat mass; *Sod1*^-/-^ mice also display reduced bodyweight and decreased visceral fat (Wang et al., 2011; Kurahashi et al., 2012; Lee et al., 2018). *hSOD1^A4V^* mice presented with an impaired early response to glucose; disturbed glucose metabolism has also been observed in *Sod1*^-/-^ mice (Wang et al., 2011, Muscogiuri et al., 2013).

Transcriptomic and metabolomic analysis of blood plasma, SC, and TA highlighted a significant dysregulation of genes and metabolites involved in glycolysis, the TCA cycle, electron transport chain, and lipid metabolism. *Sod1^-/-^*mice have altered glycolysis and gluconeogenesis and disrupted lipid metabolism, with altered hepatic and plasma lipid profiles and altered lipogenesis (Wang et al., 2012; Uchiyama et al., 2006; Kurahashi et al., 2015). However, several SOD1-ALS mouse models, without loss of SOD1 enzymatic activity, display metabolic phenotypes including reduced bodyweight, reduced fat mass, dyslipidaemia, and reduced glucose utilisation (Maksimovic et al., 2023), suggesting gain of function could also be an underlying cause.

The alterations identified in our *hSOD1^A4V^* mice align with growing evidence implicating metabolic dysfunction as a critical feature of ALS pathogenesis. Recently, neuronal polyunsaturated fatty acids have been shown to have a protective effect in *Drosophila* of ALS expressing the pathological *C9orf72* repeat expansion (Giblin et al., 2025). The polyunsaturated fatty acids found to be downregulated in affected flies overlap (with the same direction of change) with those detected in our *hSOD1^A4V^*mice, including eicopentaeonic acid, palmitoleic acid, palmitic acid and oleic acid.

Giblin *et al*., also highlight reduced expression of the genes fatty acid synthase (*FASN*) and very long chain fatty acid elongase 6 (*ELOVL6*) in C9orf72-ALS *Drosophila* and postmortem spinal cord from C9orf72-ALS patients (Giblin et al., 2025). This is consistent with data from our *hSOD1^A4V^* mice in which *Fasn* is significantly downregulated in the SC of 20-month-old *hSOD1^WT/A4V^*(p 0.044) and the TA of 12-month-old *hSOD1^WT/A4V^* (p 0.047) animals and expression of *Elovl6* is significantly reduced in the SC of 12-month-old *hSOD1^WT/A4V^* mice (p 0.045). The prospect of a shared metabolic signature involving fatty acid changes across different ALS models with various mutations is key area for future research with the potential to develop treatments applicable to all ALS patients, regardless of the aetiology of their disease. Previous studies in ALS patients and transgenic mouse models have demonstrated perturbations in glucose metabolism, mitochondrial dysfunction, and energy deficits (Bame et al., 2014; Fernández-Beltrán et al., 2021; Lee et al., 2021). The increased pyruvic acid and TCA intermediates and dysregulated lipid metabolism in our model suggest a shift towards alternative metabolic pathways, possibly as a compensatory response to impaired energy homeostasis. Moreover, the upregulation of genes involved in mitochondrial electron transport chain complexes in *hSOD1^WT/A4V^* animals suggests an attempt to counteract mitochondrial dysfunction, a well-established feature of ALS (Kiskinis et al., 2014; De Giorgio et al., 2019).

Despite the lack of overt neurodegeneration, our findings suggest that metabolic dysregulation may precede the clinical manifestation of ALS. This aligns with the concept of a pre-symptomatic metabolic shift, where bioenergetic changes occur before significant neuromuscular impairment (Xia et al., 2023; Dorst et al., 2023). The identification of altered pathways related to glycolysis, lipid metabolism, and oxidative phosphorylation raises the possibility that metabolic biomarkers could serve as early indicators of ALS risk or disease progression.

The absence of neurodegeneration in our model reinforces the need for additional triggers beyond SOD1 mutation alone (Al-Chalabi & Hardiman, 2013). Time is likely an important factor, whereby physiological levels of SOD1 may require longer than the lifespan of the mouse to reach a threshold of disordered SOD1 needed to manifest clinical phenotypes. We also note that the animals in this study had a single genetic background and thus provided only a snapshot of possible genetic interactions. Future studies should explore whether aging, environmental factors, or seeding paradigms induce more advanced phenotypes in these mice. Exposure of our humanized *hSOD1^A4V^* mice to an oxidant diet may exacerbate or expose further ALS-like phenotypes due to the paucity of SOD1 enzymatic activity (Eom et al., 2022). Additionally, targeting the metabolic pathways identified in our study, such as glycolysis and lipid metabolism, may provide novel therapeutic strategies for delaying ALS onset in genetically predisposed individuals (Bame et al., 2014; Fernández-Beltrán et al., 2021). Our findings highlight the importance of preclinical ALS models that mimic the clinically silent phase of disease, allowing for biomarker discovery and new mechanistic insights into ALS aetiology. The presence of the human *SOD1* gene at the endogenous locus also offers a platform to test therapeutics that target the human gene or gene products in a physiological context.

The observed metabolic and transcriptomic changes suggest that early dysfunctions in energy metabolism may precede motor impairment, aligning with clinical observations in pre-symptomatic ALS patients. While our model is focussed on the early stages of ALS, transgenic overexpression mice remain valuable for studying later disease stages characterized by aggressive neurodegeneration (Reaume et al., 1996; Fisher & Bannerman, 2019). Integrating these different models will provide a more comprehensive understanding of ALS pathogenesis and facilitate the development of interventions tailored to specific disease phases. Our humanised *hSOD1^WT^* and *hSOD1^A4V^*mouse strains are freely available from EMMA with the IDs EM:13075 and EM:14880, respectively.

## MATERIALS AND METHODS

### Animals

The generation of humanised *hSOD1^WT^* mice has been previously described (Devoy et al., 2021). To generate humanised *hSOD1^A4V^* mice, the A4V point mutation (C->T at g.130) was introduced into the humanised *SOD1* allele in *hSOD1^WT^*zygotes via CRISPR/Cas9 editing. A single-stranded oligodeoxynucleotide (ssODN) donor was used as a repair template. The CRISPR/Cas9 system components were delivered as ribonucleoprotein complexes into 1-cell stage *hSOD1^WT^*embryos by electroporation. The sequence of the sgRNA used in the generation of the *hSOD1^A4V^*line was: 5′-GTCGCCCTTCAGCACGCACACGG-3′. The sequence of the ssODN donor was 5′-TTCCTGCGGCGCCTTCCGTCCGTCGGCTTCTCGTCTTGCTCTCTCTGGTCCCTCCGGAGGAGGCCGCCGCGCG TCTCCCGGGGAAGCATGGCGACGAAGG**<u>T</u>**CGTGTGCGTGCTGAAGGGCGACGGCCCAGTGCAGGGCATCAT CAATTTCGAGCAGAAGGCAAGGGCTGGGACGGAGGCTTGTTTGCGAGGCCGCTCCCA-3′ (mismatch to introduce C->T point mutation underlined). Electroporated embryos were re-implanted into CD1 pseudo-pregnant females. F_0_ progeny were backcrossed for one generation into C57BL6/J to obtain germline transmission, and the resulting F_1_ offspring were screened for successful homology-directed repair (HDR) and integration of the A4V mutation by PCR followed by Sanger sequencing (Table 1). To exclude off-target integration of the donor ssODN, copy counting of the donor sequence was carried out using ddPCR (Table 1). Following confirmation of successful targeting, founder *hSOD1^A4V^*mice were crossed to *hSOD1^WT^* mice to obtain compound heterozygotes (*hSOD1^WT^*/ *hSOD1^A4V^*), which were then intercrossed to produce experimental animals.

**Table 1.**
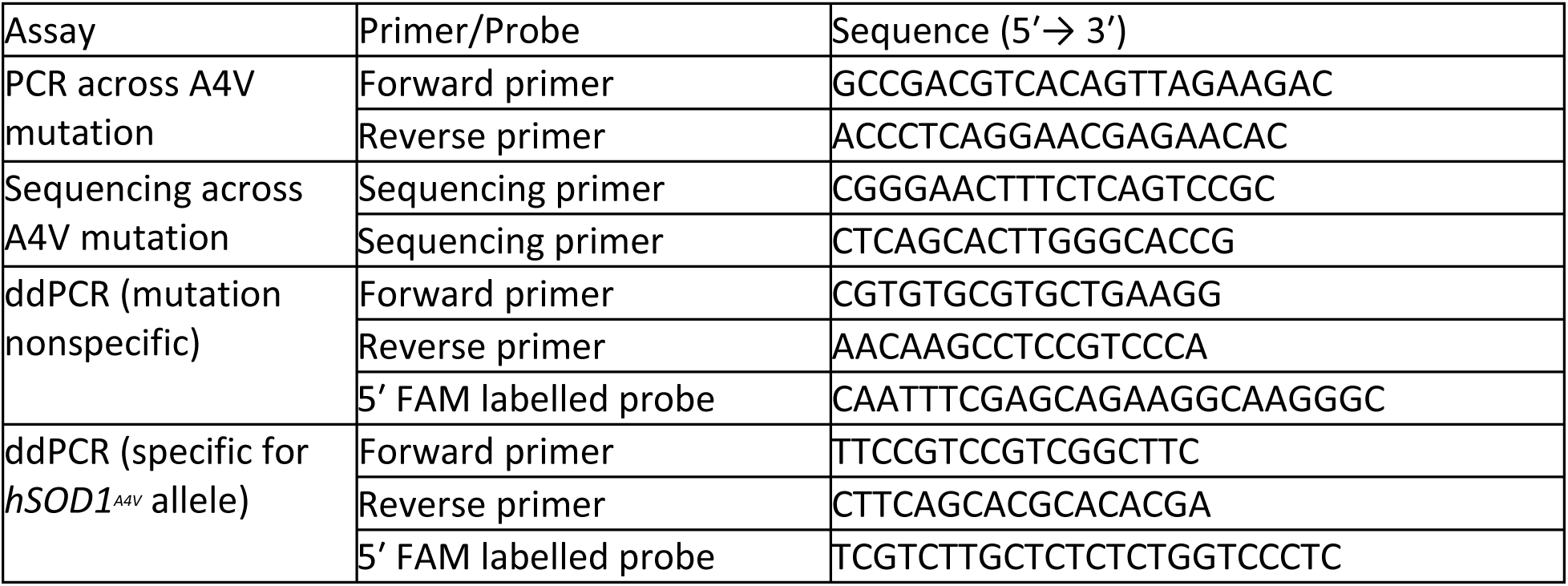
PCR primers, sequencing primers, and probes for screening of *hSOD1^A4V^* mice.

**Table 2.**
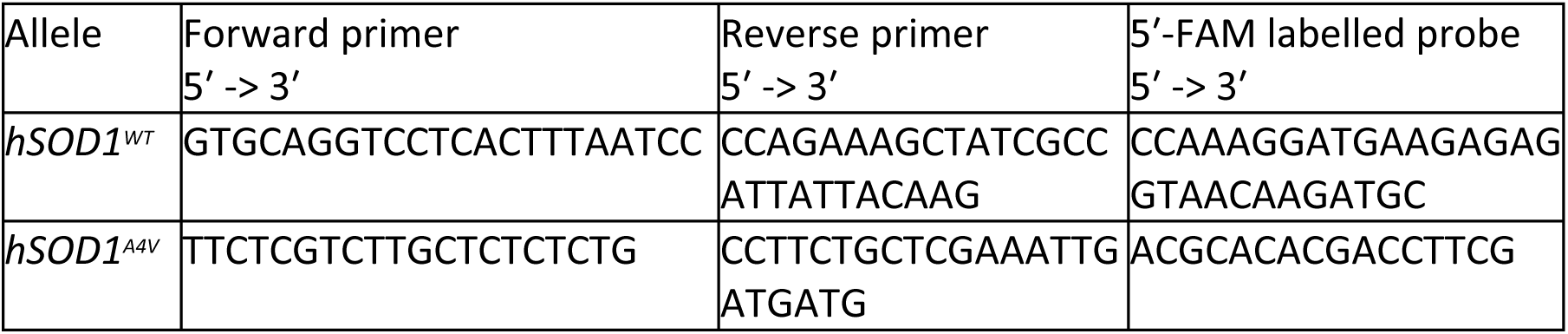
Genotyping primers and probes.

Experimental animals were considered to be on a mixed 129/SvJ x C57BL/6J background, being generation N4 on C57BL/6J (the 129/SvJ contribution is derived from the embryonic stem cells used for gene targeting to create the background *hSOD1^WT^*strain).

Mice were housed in standard cages of up to 5 animals, with *ad libitum* access to food and water, a 12h:12h light cycle with lights-on at 7am, and environmental temperature and humidity at 21±2°C and 55±10%, respectively. All animal husbandry and experiments were performed at the MRC Harwell Institute and were in accordance with the Animals (Scientific Procedures) Act 1986, Home Office guidelines and the local ethics guidelines issued by the Medical Research Council. All reasonable measures were taken to minimise animal distress and suffering.

### Genotyping

Animal genotyping was performed on genomic DNA extracted from ear clip biopsies using Loss-Of-Allele copy number qPCR with the primers and probes:

### Experimental groups

Longitudinal behavioural, motor, and metabolic phenotyping were performed at regular intervals on both male and female animals from two independent cohorts. Data from the two cohorts were pooled. Tissues were harvested from these two cohorts for histology and molecular phenotyping at 24 months old. Tissues were harvested from additional cohorts for histology and molecular phenotyping at earlier timepoints.

In all experiments, *hSOD1^A4V^*mice were compared to *hSOD1^WT/WT^* littermate controls, which have a genomically humanised *SOD1* locus but lack the A4V mutation.

### Motor Phenotyping

All phenotyping was performed in the light phase (7am-7pm). For all tests, mice were acclimatised to the testing room for at least 30 minutes prior to starting the procedure.

At each time point (6-, 12-, 18-and 24-months-old) motor and behavioural phenotyping tests were performed in the following order: open field (data not shown), combined SHIRPA (Rogers et al., 2001) and dysmorphology (data not shown), gait analysis, Locotronic (data not shown), Rotarod, grip strength, automated wheel running (data not shown).

### Accelerating Rotarod test

The accelerating Rotarod test (4–40 r.p.m. over 5 min; Ugo Basile) consisted of 6 trials performed on three days per week over two consecutive weeks. Mice were given one trial per day. The first 4 trials were considered training and no results were recorded. From the final two trials, the longest trial time was recorded. The end of the trial was classified as the time when the mouse fell from the rotarod or made three consecutive passive rotations. The maximum trial duration was 5 minutes.

### Grip strength

Grip strength was measured using a grip strength meter (Bioseb). To measure forelimb grip, mice were lifted gently by the base of the tail and held with their forelimbs in contact with the grid of the grip strength meter. Mice were then pulled horizontally away from the mesh in one smooth motion without excessive force and the resulting grip strength value was recorded in grams. To measure forelimb plus hindlimb grip the same procedure was applied, however, mice were now allowed to grip the grid with all four limbs. Three measurements of forelimb grip and three measurements of forelimb plus hindlimb grip were collected from each mouse at each time point.

To minimise interoperator variability, at each phenotyping time point the same researcher performed all grip strength measurements. For forelimb, and forelimb plus hindlimb grip strength the median measurement from each animal was divided by the weight of the animal and the resulting value was used for analysis.

### Gait analysis

The MouseWalker system is used to examine gait in freely walking rodents, similar to the footprint test. The MouseWalker apparatus is described in Mendes et al., 2015 and consists of an illuminated acrylic glass catwalk with a high-speed camera to record mouse foot placement.

The MouseWalker procedure was performed in the dark. Mice were placed at the end of the catwalk and allowed to walk freely along it for 1 minute. Videos captured by the camera were analysed frame by frame to quantify gait parameters.

### Metabolic Phenotyping

#### Echo MRI

Echo MRI measures body composition in conscious animals; providing measurements of whole-body fat mass, lean mass and water content. Echo MRI was performed using a mouse Echo MRI machine (Echo Medical Systems).

#### Intra-peritoneal glucose tolerance testing (IPGTT)

Mice were fasted overnight prior to the test. EMLA local anaesthetic cream was applied to the tail 30 minutes prior to the test. Blood was sampled from the lateral tail vein and glucose measurements performed using an AlphaTrak meter and AlphaTrak test strips. For the T0 baseline measurement, blood glucose was measured prior to glucose administration. Glucose solution was then administered via IP injection at a dose of 2 g/kg. Blood glucose was measured 15, 30, 60 and 120 minutes after glucose administration. Statistical analysis was performed on data from the individual sampling time points and on the area under the curve (AUC) across all time points.

#### Blood sampling for glutathione measurement and metabolomics

Mice were fasted for approximately 15 hours before blood sampling from the lateral tail vein. EMLA cream was applied to the tail 30 minutes prior to blood sampling, then thoroughly removed with an ethanol wipe before a small incision was made at the base of the tail. Between 100ul and 200ul blood was collected into heparin-coated tubes. The volume of extracted blood did not exceed 15% total blood volume. 50ul whole blood was immediately mixed with 50ul meta-phosphoric acid (MPA) buffer (5% w/v meta-phosphoric acid, 2mM diethylenetriaminepentaacetic acid) and deprotonated on ice for 10 minutes. This whole blood/MPA buffer mix was used for measurement of glutathione. The remaining whole blood was left to coagulate on ice for 40 minutes, then centrifuged at 2000 x g for 10 minutes at 4°C. The plasma and cell pellet were separated and both were flash frozen in liquid nitrogen and stored at-80°C before analysis.

#### Monthly weights

Mice were weighed monthly at approximately the same time of day (between 8am and 12pm). Mice that reached 10% weight loss compared to their individual maximum body weight were weighed more frequently (weekly or daily).

### Survival

Animal health and wellbeing was assessed daily. The humane end points for this study included: weight loss exceeding 20% of maximum bodyweight, piloerection or other loss of condition, postural abnormalities (e.g. hunched appearance, tremors or ataxia impinging on the ability to feed), abnormal behaviour impinging on the welfare of the animal (e.g. lethargy, extreme hyperactivity), seizures, and age-related loss of condition. Mice culled for “non-ALS” causes (e.g. fighting, tumours) were not included in the survival analysis. Mice that did not develop ALS symptoms were culled by 27 months of age. Survival data were analysed by Kaplan-Meier.

Statistical significance was determined using a log-rank (Mantel-Cox) test where P < 0.05 was considered significant.

### Harvesting of tissues

Animals were terminally anesthetised with 1000 mg/kg of pentobarbitone administered via i.p. injection. Once sufficiently anesthetised all, or a subset, of the following tissues were harvested from each animal: brain, SC, quadriceps, TA, EDL, soleus, kidney, liver, lumbrical muscles, and tail. Further processing of fresh tissue differed depending on the experiment.

For protein analysis, tissues were flash frozen in liquid nitrogen and stored at-80°C until use. For RNA analysis, tissues were immediately immersed in ice-cold RNAlater solution (Sigma Aldrich) and stored at-20°C in RNAlater until use. For muscle fibre size measurement and ATPase staining, TA muscles were attached to cork using trigacanth gum, covered in OCT and flash frozen in isopentane over liquid nitrogen. Frozen muscles were stored at-80°C until use. For motor neuron counting, lumbar spinal cords were embedded in OCT and frozen in isopentane over dry ice. Embedded spinal cords were stored at-80°C until use. For the analysis of neuromuscular junctions (NMJs), hindlimb lumbrical muscles were extracted, fixed for 10 minutes in 4% PFA and stored in PBS until staining.

TA and soleus muscles for NMJ analysis were extracted, pinned out, and fixed in 4% PFA for between 20 minutes and 3 hours. Fixed TA and soleus muscles were stored in PBS for up to one month before further processing.

### Histology, immunohistochemistry and immunofluorescence

#### MN counts

Our protocol for motor neuron counting has been published (Weydert & Cullen, 2010). Briefly, fresh-frozen lumbar spinal cords were embedded in OCT and sectioned using a cryostat. 20 µm thick transverse sections were cut spanning the L2-L6 region of the lumbar spinal cord and mounted on glass slides. The sections were divided into 3 series such that the distance between sections within each series was 60µm. Sectioning was performed by histology staff at the MRC Harwell institute.

To visualise motor neurons, sections from *hSOD1^WT^* mice were stained with gallocyanin and sections from *hSOD1^A4V^* mice were stained with cresyl violet. Stained sections were mounted and imaged on a Nanozoomer slide scanner (Hamamatsu). For each animal, spinal motor neurons in the sciatic pool of the lumbar spinal cord were counted in a minimum of 40 sections spanning the L2-L6 region of the spinal cord. Spinal cord regions L2-L6 were identified from the spinal cord morphology. Motor neurons were differentiated from other cell types by their large size, dense staining pattern and the presence of multiple dendrites. Motor neuron counting was performed in the NDP.view 2 software (Hamamatsu). Analyses are presented as the average number of sciatic motor neurons per section in the L3-L5 region of the spinal cord.

#### NMJ analysis on lumbrical muscles

The protocol for staining and analysis of NMJs in lumbrical muscles was adapted from Sleigh et al., 2014. Fixed hindlimb lumbrical muscles in 24 well plates were washed once in PBS for 30 minutes. Muscles were then permeabilised in 2% triton X-100 (v/v) for 30 minutes, followed by a 30-minute incubation in blocking solution (4% BSA (w/v), 1% triton X-100 (v/v), PBS), and incubation in primary antibodies against neurofilament (2H3; 1:50) and synaptophysin (SV2; 1:100) in blocking solution at 4°C overnight. Muscles underwent 3 x 30-minute washes in PBS, followed by incubation with anti-mouse IgG (1:500, Alexa Fluor 488, ThermoFisher) and α-Bungarotoxin (α-BTX; 1.5 µg/ml; tetramethylrhodamine; Biotium) in PBS for 2 hours at room temperature. Muscles were washed 3 x 30-minutes in PBS and mounted on glass slides in ProLong Gold mounting media (ThermoFisher).

Samples were imaged at 488 nm and 555 nm wavelengths on an LSM 700 confocal microscope at 20x magnification. NMJ analysis on Z-stack maximum intensity projections was performed in ImageJ. NMJs were recorded as innervated or denervated; partial denervation was not assessed due to high non-specific staining with antibodies against 2H3 and SV2 in older animals. At least 40, but usually 100-200 NMJs were counted from each animal.

#### NMJ analysis on TA and soleus muscles

The TA and soleus muscles from 6-month-old *hSOD1^A4V^* animals were injected with tetanus toxin heavy chain (atoxic fragment of tetanus neurotoxin, H_C_T) as part of axonal transport experiments; H_C_T labels neuronal endosomes and hence stains the axon/presynapse without the need for antibodies. The TA and soleus muscles from 24-month-old mice were labelled using antibodies against SV2/2H3.

The protocol for processing of TA and soleus muscles for NMJ analysis was adapted from Bolatto et al., 2021. TA and soleus muscles were fixed for 30-180 minutes in 4% PFA, teased apart into bundles of approximately 5-20 fibres and transferred to PBS in 24 well plates. Fibre bundles were washed once in PBS for 20 minutes. Fibre bundles were then stained with α-BTX (1:500, Alexa Fluor 488 or tetramethylrhodamine) for 1 hour at room temperature. Fibre bundles from 24-month-old animals were permeabilised in 2% Triton X-100 (v/v) for 90 minutes, blocked in 4% BSA (w/v) and 1% Triton X-100 (v/v) for 30 minutes and incubated in primary antibodies against 2H3 (1:50) and SV2 (1:100) in blocking solution at 4°C overnight. Fibre bundles from 24-month-old animals were washed 4 x 20 minutes in PBS before incubation in anti-mouse IgG (1:250, Alexa Fluor 488, ThermoFisher) for 1 hour at room temperature. All fibre bundles were then washed 4 x 20 minutes in PBS and mounted on glass slides in ProLong Gold mounting media (ThermoFisher).

Samples were imaged at 488 nm and 555 nm wavelengths on an LSM 700 confocal microscope at 20x magnification. NMJ analysis on Z-stack maximum intensity projections was performed in ImageJ. NMJs were recorded as innervated or denervated; partial denervation was not assessed due to poor quality of staining of the postsynaptic NMJ with α-BTX. At least 40 NMJs were counted from each muscle.

### *In vivo* axonal transport

#### Intramuscular injections of H_C_T

Fluorescently labelled atoxic fragment of tetanus neurotoxin (H_C_T) was prepared (H_C_T residues 875– 1315 fused to a cysteine-rich region and a human influenza haemagglutinin epitope), and subsequently labelled with Alexa Fluor 555 C2 maleimide (Thermo Fisher Scientific, A-20346), as previously described (Sleigh et al., 2020; Tosolini et al., 2021). Mice were anesthetized using isoflurane, the fur on lower leg was shaved, and mice were placed on a heat-pad ready for intramuscular injections. Upon the absence of reflexes, a small incision was made above the upper tibialis anterior muscle (TA) of both hindlimbs. Guided by previously established motor end plate maps (Mohan et al., 2014), injections were made into both the left and right TA with 7.5–10 mg of H_C_T in PBS in a volume of ∼3.5 µL using pulled graduated, glass micropipettes (Drummond Scientific, 5-000-1001-X10), as previously described (Mohan et al., 2015). When complete, the skin was sutured, and mice were monitored for up to 1 h before returning to the home cage.

#### In vivo imaging of signalling endosomes

To visualise signalling endosome transport *in vivo,* time-lapse confocal microscopy of motile H_C_T-containing signalling endosomes in TA-innervating motor axons was performed, as described previously (Sleigh et al., 2020; Tosolini et al., 2021). At least 4 h after H_C_T intramuscular injections, mice were re-anaesthetised with isoflurane, and the sciatic nerves were exposed with the removal of overlying skin, musculature and connective tissue, and a small piece of parafilm was inserted to separate the sciatic nerve from the underlying connective tissue/musculature. The mouse and anaesthetic nosepiece were then transferred to a customised stage on an inverted LSM780 confocal microscope (Zeiss) enclosed within an environmental chamber maintained at 37^0^C. Time-lapse microscopy was performed using a 40x, 1.3 NA DIC Plan-Apochromat oil-immersion objective (Zeiss) focusing on thicker axons using an 80x digital zoom (1024 x 1024, <1% laser power). Only the axons that contained H_C_T-containing signalling endosomes were selected for signalling endosome transport, with areas containing the node of Ranvier purposely avoided (Tosolini et al., 2024).

Frame intervals of 1.0 – 1.3 s were used when acquiring transport videos of motile H_C_T-positive signalling endosomes. When the first leg was complete, the same process was repeated for the contralateral limb. All imaging was concluded within a maximum of 90 minutes from initiating re-anaesthesia.

#### Tracking analysis

Confocal ‘‘.czi’’ images were opened in FIJI (http://rsb.info.nih.gov/ij/), and converted to ‘‘.tiff’’. Using the TrackMate plugin (Ershov et al., 2022), H_C_T-positive signalling endosomes were tracked manually (Tosolini et al., 2022, 2024), using the following criteria: 1) organelles moving for >10 consecutive frames were included, while terminal pauses, which we defined as the absence of movement in >10 consecutive frames, were excluded. In addition, tracking never commenced on an immotile organelle; 2) a minimum of 20 signalling endosomes were selected per axon; 3) at least 3 individual axons were assessed per limb. A pause was defined as a previously motile organelle with a velocity of <0.1 mm/s between consecutive frames (to account for a potential breathing/vasculature artifact), and the time paused (%) was defined by dividing the number of pauses by the total number of frame-to-frame movements assessed from an individual axon.

### Muscle fibre typing

Muscle ATPase staining was performed to differentiate type I, II and IIC muscle fibres. Flash frozen TA muscles were cut into 12 μm thick transverse sections on a cryostat. Sections were preincubated in sodium barbital solution at pH 4.5 (35 mM sodium barbital, 60 mM sodium acetate, HCl) or pH 10.8 (20 mM sodium barbital, 36mM CaCl_2_, NaOH). Sections were rinsed once in distilled water followed by incubation in ATP solution (20 mM sodium barbital, 9 mM CaCl_2_, 2.7 mM ATP; pH 9.5). Sections were then washed 3 x 3 minutes in 1% CaCl_2_ (w/v) before incubation for 10 minutes in 2% CoCl_2_ (w/v). Sections were washed in 5 changes of 5 mM sodium barbital. Sections were then incubated in 1% ammonium sulphide (v/v) for 30 seconds, washed in 6 changes of water and mounted using Aquamount (Life Technologies). Stained sections were digitalised on a Nanozoomer slide scanner (Hamamatsu). Muscle fibre type was assessed in a 750 µm x 750 µm area positioned at approximately the same location within each section. For each animal muscle fibre type was assessed at pH 4.5 and pH 10.8 on an average of 2 sections spaced 400 µm apart.

### Determination of muscle fibre size and central nucleation

Fresh frozen TA muscles were cut into 8 μm thick transverse sections on a cryostat. Sections were transferred to glass slides, stained with Haemotoxylin and Eosin (H&E), and digitized using a slide scanner (Zeiss Axio Scan.Z1). For each muscle, three 500µm x 500µm regions were selected for determination of muscle fibre size. Segmentation of muscle fibres was performed in Ilastik ^408^ and ImageJ (National Institutes of Health) and checked manually. The minimum Feret diameter for each fibre was calculated in ImageJ. Central nucleation was assessed manually in 500 fibres per muscle.

### Biochemical and molecular analyses

#### RNA extraction for Reverse transcription quantitative PCR (RT-qPCR) and RNA sequencing (RNA-seq)

RNA was extracted from CNS tissues using an RNeasy Lipid Tissue Mini Kit (Qiagen), from muscle using an RNeasy Fibrous Tissue Mini Kit (Qiagen), and from Kidney and liver using an RNeasy Mini Kit (Qiagen). RNA quantity was assessed using a NanoDrop Spectrophotometer (ThermoFisher). RNA quality was assessed using a 2100 Bioanalyzer (Agilent) where an RNA integrity number (RIN) of >7 was deemed acceptable for RT-qPCR and RNA-seq. Extracted RNA was stored at-80°C until use.

#### Reverse transcription quantitative PCR (RT-qPCR)

Complementary DNA (cDNA) was generated using a High-Capacity cDNA Reverse Transcription Kit (Applied Biosystems). 10 ng cDNA was used to perform RT-qPCR in triplicate, using 5 μl cDNA, 10 μl Fast SYBR Green Master Mix (Thermo Fisher) and 0.2 μM each forward and reverse primer in a total reaction volume of 20 μl. RT-qPCR was performed on a 7500 Fast Real-Time PCR System (Applied Biosystems) using the following conditions: 95 °C for 10 minutes, 40 cycles of 95 °C for 15 seconds, 60 °C for 1 minute. The sequences of primers used in this study are listed in Table 3. Relative quantification of gene expression was performed using the 2^-ΔΔ*CT*^ method. *Gapdh* was used as a housekeeping gene for normalisation of *SOD1* gene expression.

**Table 3.**
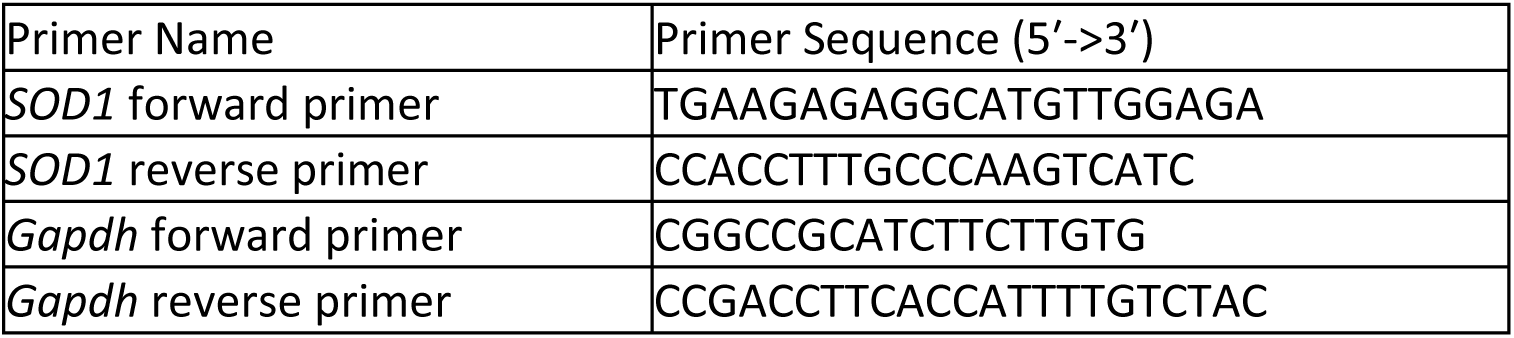
List of RT-qPCR primers.

### Protein extraction from mouse tissues

Flash-frozen tissues, stored at-80°C, were thawed on ice and homogenised in 25 volumes of ice-cold lysis buffer (PBS, Complete Protease Inhibitor Cocktail (Roche), 0.5% NP-40). Lysates were centrifuged at 18,000 x g for 30 minutes at 4°C and the supernatants were collected and stored at

−80°C until analysis. Protein concentration was measured using a DC Protein Assay (BioRad).

### Western blotting

Ten to fifteen micrograms of protein per sample was used for reducing SDS-PAGE on Novex 14% Tris-Glycine gels (ThermoFisher) followed by western blotting. Blots were probed with a pan-SOD1 antibody raised against a peptide corresponding to amino acids 131-153 of the SOD1 protein, which are identical in human and mouse SOD1, followed by IRDye 680RD conjugated secondary antibody (LI-COR) (Jonsson et al., 2004). SOD1 immunofluorescence was normalised to the total protein content of each sample, measured using a Revert 700 Total Protein Stain (LI-COR). Images were taken using an Odyssey CLx imager (LI-COR).

### In-gel dismutase activity assays

The protocol for in-gel dismutase activity assays was adapted from Weydert & Cullen, 2010. 15-30 ug of protein per sample was separated using clear native PAGE (CN-PAGE) on 3-12% NativePAGE Bis-Tris gels (ThermoFisher). Following CN-PAGE, gels were soaked in SOD1 stain solution (50 mM potassium phosphate, 336 µM nitro blue tetrazolium, 170 µM riboflavin, 21 mM TEMED) in the dark for 45 minutes. Gels were then rinsed with ddH_2_O twice and placed under a fluorescent light for approximately 1 hour, during which time clear bands corresponding to areas of SOD1 activity began to appear. Gels were then washed 3 times with ddH_2_O and left under ambient light for 18-24 hours to allow for further band development. Gels were imaged using a ChemiDoc XRS+ imaging system (BioRad) with a white light conversion filter.

### Spectrophotometric dismutase activity assays

The protocol for a high throughput spectrophotometric SOD dismutase activity assay was adapted from Peskin and Winterbourn, 2017. Protein samples were thawed on ice and dispensed into 96-well assay plates. Depending on the tissue, 1-5ug of protein per sample was used for dismutase activity assays. 200 µL of reagent mixture (100 mM pH 8.0 sodium phosphate, 100 µM DTPA, 10 mM hypoxanthine, 50 µM WST-1, 10 µg/mL catalase, 0.1 units/mL xanthine oxidase (Sigma)) was added to every well using a multichannel pipette. Plates were shaken for 30 seconds to mix then absorbance was read immediately at 438 nm wavelength every 40 seconds for 3 minutes at 25℃ SOD1 activity was determined based on comparison to 12-point standard curve of bovine SOD1 ranging from 1-1000 ng/well. 1 unit of activity was defined as the amount of bovine SOD1 required for 50% inhibition of tetrazolium reduction (approximately 25 ng).

### ELISA for soluble disordered SOD1 (misELISA)

ELISA plates (Maxisorp, Nunc) were coated overnight at room temperature with 640 ng/mL of anti-SOD1 (human) monoclonal antibody (raised against amino acids 57-72, exposed only in disordered SOD1) in 100 mM Tris-HCl. After washing (PBS and 0.05% Tween-20) and blocking, 25 μg of protein samples and apo-SOD1 standards diluted in PBS, protease inhibitor and 40 mM iodoacetamide were incubated for 90 minutes at room temperature. Each well was washed thrice before addition of 1.5 μg/ml goat antibody, raised against mouse SOD1 for 1 h at room temperature. After washing, anti-goat HRP (Invitrogen, 1:5000) was added for 1 h at room temperature. Finally, after washing, wells were incubated in TMB substrate for 15 minutes. Sulphuric acid (2M) was added to stop the reaction and the absorbance read at 450 nm.

### Transcriptomics and Metabolomics

#### Quantseq 3′ mRNA sequencing

Library preparation and Illumina sequencing were performed by Lexogen. cDNA libraries were prepared using QuantSeq 3′ mRNA-Seq Library Prep FWD kits (Lexogen) and sequenced on an Illumina platform to generate single-end reads. Basic statistics are listed in Table 4.

**Table 4.** Read counts from Quantseq 3′ mRNA sequencing runs. mo, month-old.

| Sample group | Total read count (Median) | Uniquely mapped reads (Median) | Uniquely mapped reads % (Median) | Average input read length (Median) |
| --- | --- | --- | --- | --- |
| 6-month-old spinal cord | 8200155 | 5369403 | 65.7 | 86 |
| 6-month-old TA muscle | 7379234 | 4984674 | 68.0 | 90 |
| 12-month-old spinal cord | 4860088 | 3198704 | 66.2 | 66 |
| 12-month-old TA muscle | 4767525 | 3146542 | 65.7 | 67 |
| 20-month-old spinal cord | 5150797 | 3571121 | 69.0 | 86 |
| 20-month-old TA muscle | 4609703 | 3419747 | 74.4 | 88 |

Quality control and read trimming was performed with Trimmomatic version 0.36. Trimmed reads were then aligned to the mm39 mouse reference genome with HISAT2 (Version 2.1.0).

FeatureCounts was used to count exon spanning reads. One *hSOD1^WT/WT^* sample from the 6-month-old TA muscle dataset clearly deviated from all other samples in PCA and was excluded from downstream analyses. Differential gene expression (DE) analysis was performed using DESeq2 (Version 1.40.1). Average gene expression was obtained n=4 animals in each genotype/time point) and the proportion of DEGs between groups was determined using the Benjamini and Hochberg procedure implemented in DESeq2 for correction of p values with false discovery rate (FDR). To identify genes of considered possibly biologically relevant, analysis was also conducted using a P-value of <0.01. DESeq2 output files were exported and volcano plots developed in GraphPad Prism 10. The UpsetR package in R Studio was used to design Vennplots. Enrichr (Chen et al., 2013; Kuleshov et al., 2016; Xie et al., 2021) was used to identify significantly enriched GO terms and infer canonical pathways regulated by the identified DEGs. Chord plots were constructed according to Tang et al., 2023.

#### Preparation of samples for metabolomics

Frozen plasma from *hSOD1^A4V^* animals were thawed at room temperature and diluted 1/10 with 90% methanol. The samples were then precipitated at-20°C for 2 hours and centrifuged for 10 minutes at 17,000 x g at 4°C. The supernatants were transferred to glass vials and dried in a SpeedVac concentrator. Quality control samples were generated by pooling a small amount of all experimental samples. Internal standards of known concentration were added to each sample.

#### LC-MS

Liquid chromatography–mass spectrometry (LC-MS) experiments were performed by the Swedish Metabolomics Centre. Samples were run on an ultra-high performance liquid chromatography-quadrupole time-of-flight mass spectrometry (UHPLC/Q-TOF-MS) instrument in both positive and negative mode. To minimise the effect of instrumental drift, samples from each time point were run in a random order. Using a targeted data processing approach, 212 metabolites were annotated in samples from 4-13-month-old animals, and 195 metabolites were annotated in samples from 20-month-old animals.

#### GC-MS

Gas chromatography–mass spectrometry (GC-MS) experiments were performed by the Swedish Metabolomics Centre. Samples were run on a gas chromatography time-of-flight mass spectrometry (GC-TOF-MS) instrument. Samples from each time point were run in a random order. Using a targeted data processing approach, 94 metabolites were annotated in samples from 4-13-month-old animals, and 82 metabolites were annotated in samples from 20-month-old animals.

#### Measurement of GSSG/GSH

Reduced glutathione (GSH) and oxidised glutathione (GSSG) levels were measured in whole blood from *hSOD1^A4V^* animals using LC-MS. Before GSH/GSSG analysis, 25 µl of each sample was mixed with 75 µl extraction solution (1.67% MPA, 3.33 µM GSH as internal standard). GSH and GSSG were annotated in the MS data using a targeted processing approach.

#### Metabolomics data analysis

Data pre-processing and annotation of metabolites was performed by the Swedish Metabolomics Centre. The input for statistical analyses was a data matrix of annotated compound names with associated peak (mz/rt) intensities. Univariate analyses were performed to identify significantly differences between the levels of metabolites in *hSOD1^WT/WT^*, *hSOD1^WT/A4V^*, and *hSOD1^A4V/A4V^* animals (p > 0.05).

#### Integrated transcriptomic and metabolomic analysis

Integrated network analysis and visualization of transcriptomic and metabolic data were performed using a combination of STRING, Reactome, and BioRender. This approach allowed for the exploration of relationships between DEGs and significantly differentially expressed metabolites (DEMs) and associated metabolic pathways, providing a comprehensive view of the biological processes affected. DEGs were uploaded to the STRING database within Cytoscape (STRING v.11, Cytoscape v.3.10.2). STRING was used to predict protein-protein interactions (PPIs) based on various evidence sources, including experimental data, computational predictions, and text mining. The confidence score for PPIs was set to 0.4 to retain only interactions with sufficient reliability. The STRING network was queried against Reactome pathways to identify over-represented pathways associated with the DEGs and DEMs (Reactome v.91; Milacic et al., 2024). Enriched pathways were visualized using BioRender to create a schematic representation of the key pathways and interactions.

### Statistics

Statistical analyses were performed in GraphPad Prism (Dotmatics). Statistical significance for phenotyping and molecular experiments was determined using ANOVA (one-way, two-way, and two-way with repeated measures) or mixed effects analysis, using a significance threshold of P ≤ 0.05. Statistical analysis of RNA-seq and metabolomics experiments are described in detail in the relevant methods sections.

## ACKNOWLEDGEMENTS

The authors thank the Swedish Metabolomics Centre, Umeå, Sweden (www.swedishmetabolomicscentre.se) for metabolic profiling by GC-MS and LC-MS. We thank all staff involved at the Mary Lyon Centre at MRC Harwell, including Lydia Teboul and the Molecular and Cellular Biology Team for zygote gene targeting and genotyping; Jackie Harrison, Gemma Atkins, Dr Anju Paudyal, and Emily Ireson for animal husbandry and colony management; Dr Michelle Stewart for colony management, Adele Austin for histology support; and Clare Norris and Sarah Taylor for phenotyping support. We also thank Dr Clemens Falker-Gieske at the University of Göttingen for sharing his time and expertise to train Chloe Williams in the analysis of transcriptomic data sets.

## DATA AND RESOURCE AVAILABILITY

The official allele designation of the humanised *hSOD1^A4V^* allele is Sod1^em1(SOD1)Emcf^ (MGI:6856511). The official strain name of the humanised *hSOD1^A4V^* mice on a C57BL/6J background is B6.129-Sod1^em1(SOD1)Emcf^/H (MGI:6856512). This line is freely available from the European Mouse Mutant Archive (number to be added).

Data used for analysis are attached in the following supplementary tables:

- Supplementary Table 1 - significantly differently expressed genes in the tibialis anterior and spinal cord of *hSOD1^WT/A4V^* and *hSOD1^A4V/A4V^*animals compared to littermate controls (padj <0.1)
- Supplementary Table 2 - overlap between datasets outputted from VennPlots
- Supplementary Table 3 - top ten most upregulated and downregulated genes detected
- Supplementary Table 4 - all differentially expressed genes in the tibialis anterior of *hSOD1^WT/A4V^* and *hSOD1^A4V/A4V^* animals compared to littermate controls
- Supplementary Table 5 - all differentially expressed genes in the spinal cord of *hSOD1^WT/A4V^* and *hSOD1^A4V/A4V^* animals compared to littermate controls

## AUTHOR CONTRIBUTIONS STATEMENT

Conceptualization: E.M.C.F., T.C., J.G.; Methodology: D.T., C.W., J.G., A.P.T., G.S.; Software: C.W.; Formal analysis: C.W., D.T.; Investigation: D.T., C.W. A.P.T; Data Curation: D.T., C.W.; Writing – original draft: C.W.; Writing – review & editing: D.T., E.M.C.F., T.C., J.G, A.P.T, G.S; Visualisation: C.W., D.T., A.P.T; Supervision: E.M.C.F., T.C., J.G.; Project administration: E.M.C.F., T.C.; Funding acquisition: E.M.C.F., T.C., J.G, A.P.T, G.S.

## SUPPLEMENTARY FIGURES

**Supplementary Figure 1.**
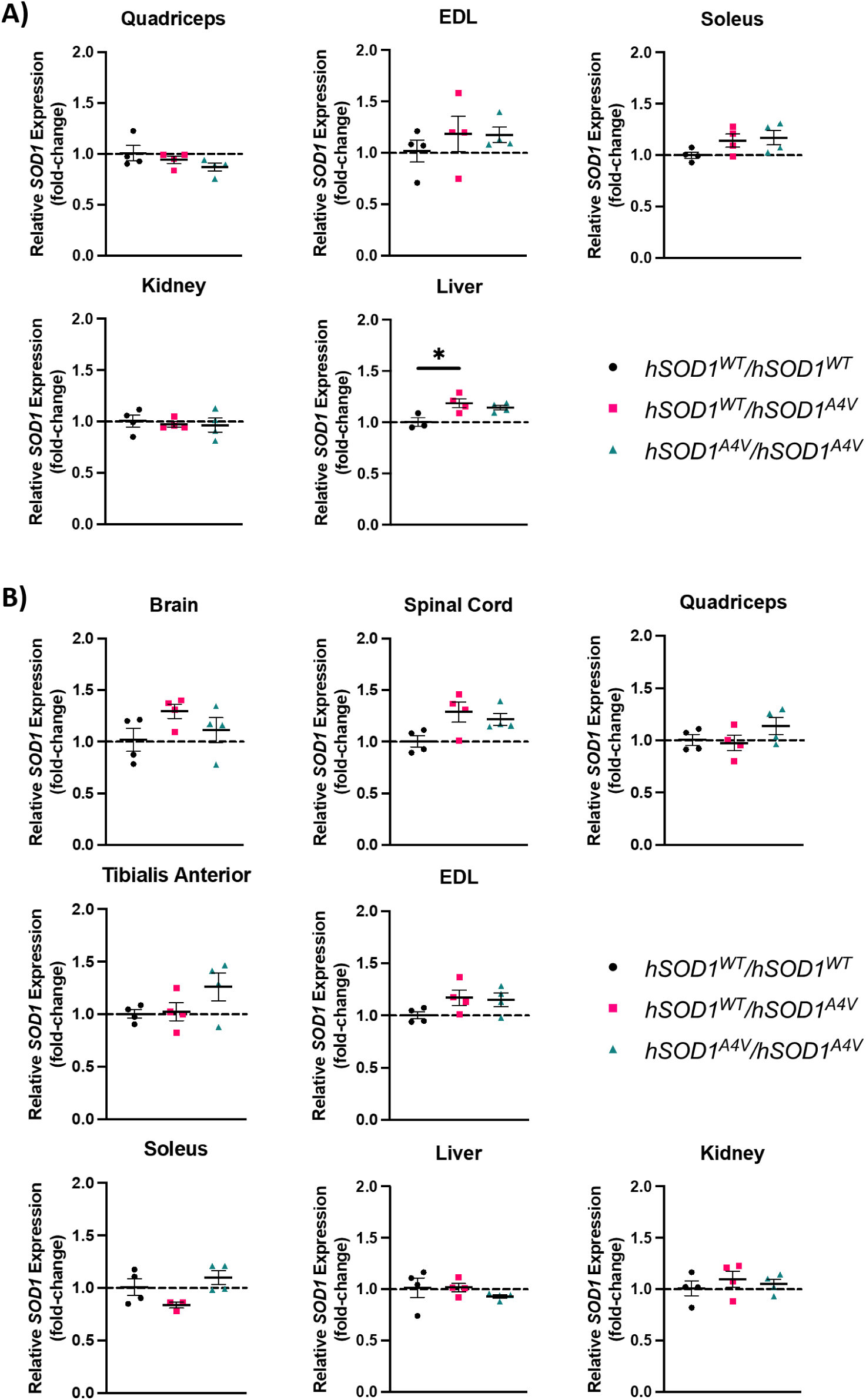
*SOD1* mRNA expression in *hSOD1^A4V^* mice. Expression of total *SOD1* mRNA using conserved mouse-human *SOD1* primers in tissues from 3-month-old A) male and B) female animals (n=3-4 animals per genotype per tissue). All values are expressed as a fold change relative to the average *SOD1* expression in *hSOD1^WT/WT^*control animals. Mean ± SEM. One-way ANOVA with Tukey’s post hoc test. \**P* < 0.05.

**Supplementary Figure 2.**
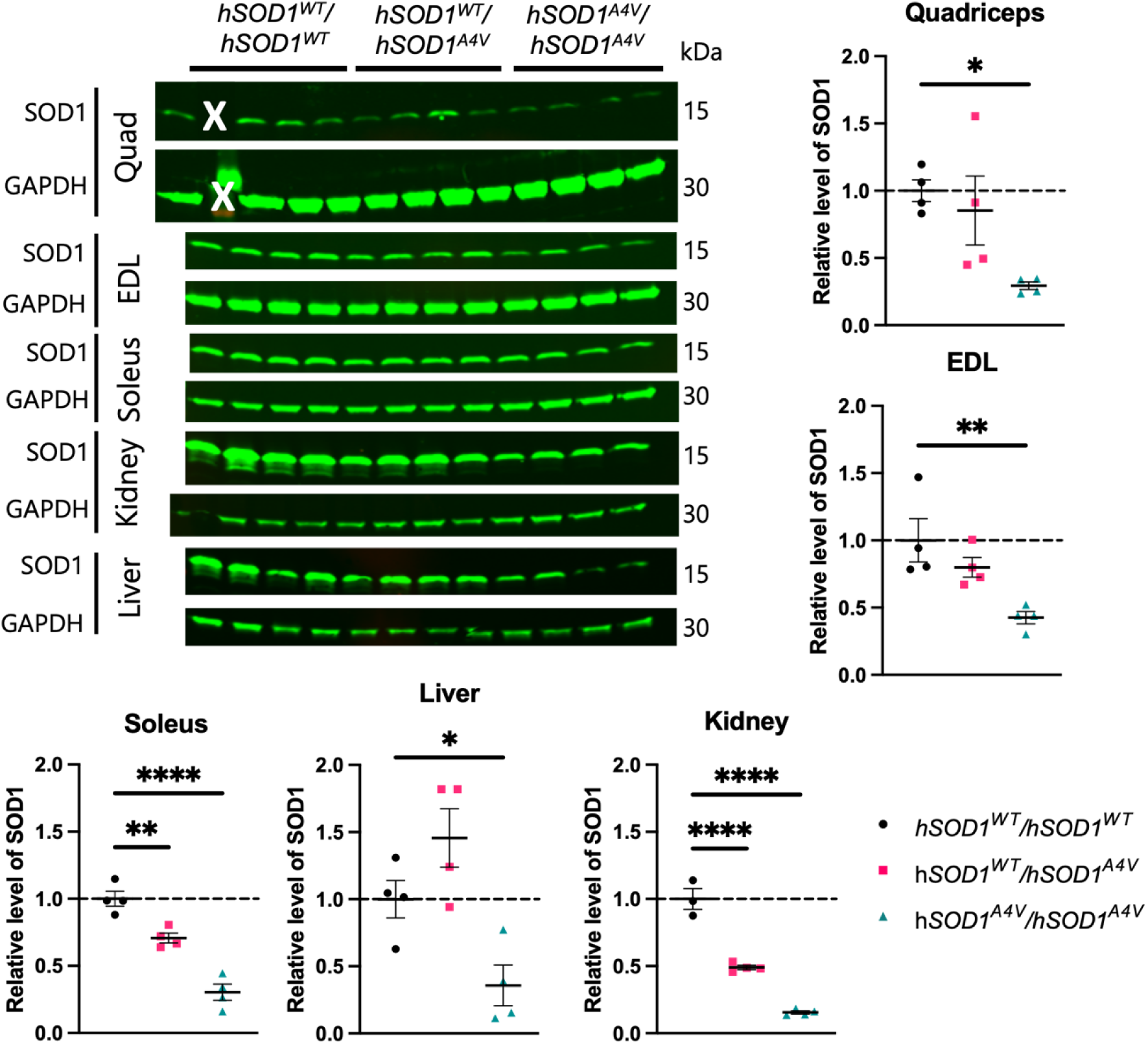
SOD1 protein male *hSOD1^A4V^* mice. Western blots using a pan-mouse-human SOD1 antibody (aa131-153) in tissues from 3-month-old male animals. Human SOD1 has lower mobility than mouse SOD1 in SDS-PAGE and appears as a higher band. Blots normalized to GAPDH. All values are expressed as a fold change relative to the average SOD1 level in *hSOD1^WT/WT^* control animals. n=4 per genotype per tissue. Mean ± SEM. One-way ANOVA with Tukey’s post hoc test. \**P* < 0.05, \*\**P* < 0.01, \*\*\**P* < 0.001, \*\*\*\**P* < 0.0001.

**Supplementary Figure 3.**
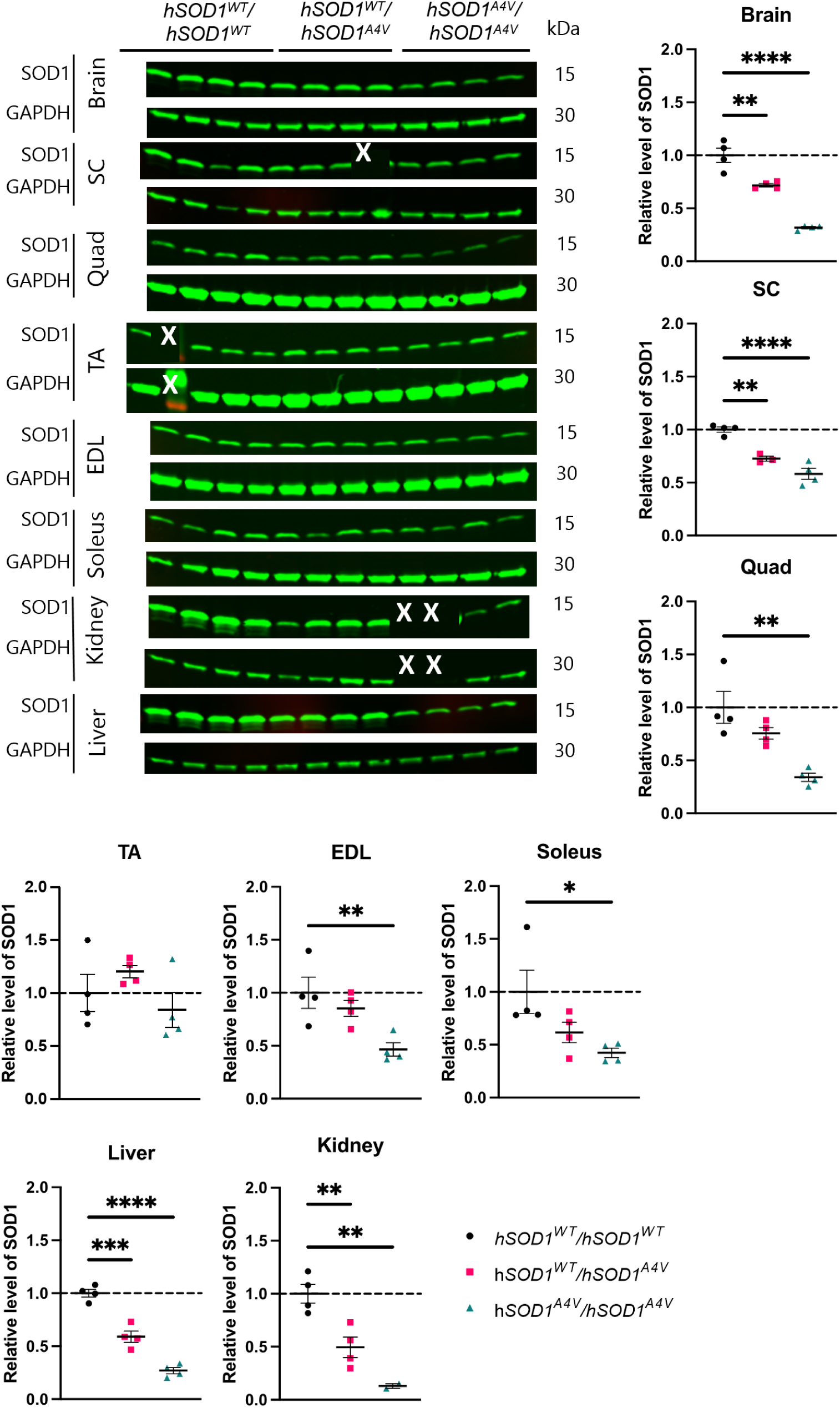
SOD1 protein female *hSOD1^A4V^* mice. Western blots using a pan-mouse-human SOD1 antibody (aa131-153) in tissues from 3-month-old female animals. Human SOD1 has lower mobility than mouse SOD1 in SDS-PAGE and appears as a higher band. Blots normalized to GAPDH. All values are expressed as a fold change relative to the average SOD1 level in *hSOD1^WT/WT^* control animals. n=4 per genotype per tissue. Mean ± SEM. One-way ANOVA with Tukey’s post hoc test. \**P* < 0.05, \*\**P* < 0.01, \*\*\**P* < 0.001, \*\*\*\**P* < 0.0001.

**Supplementary Figure 4.**
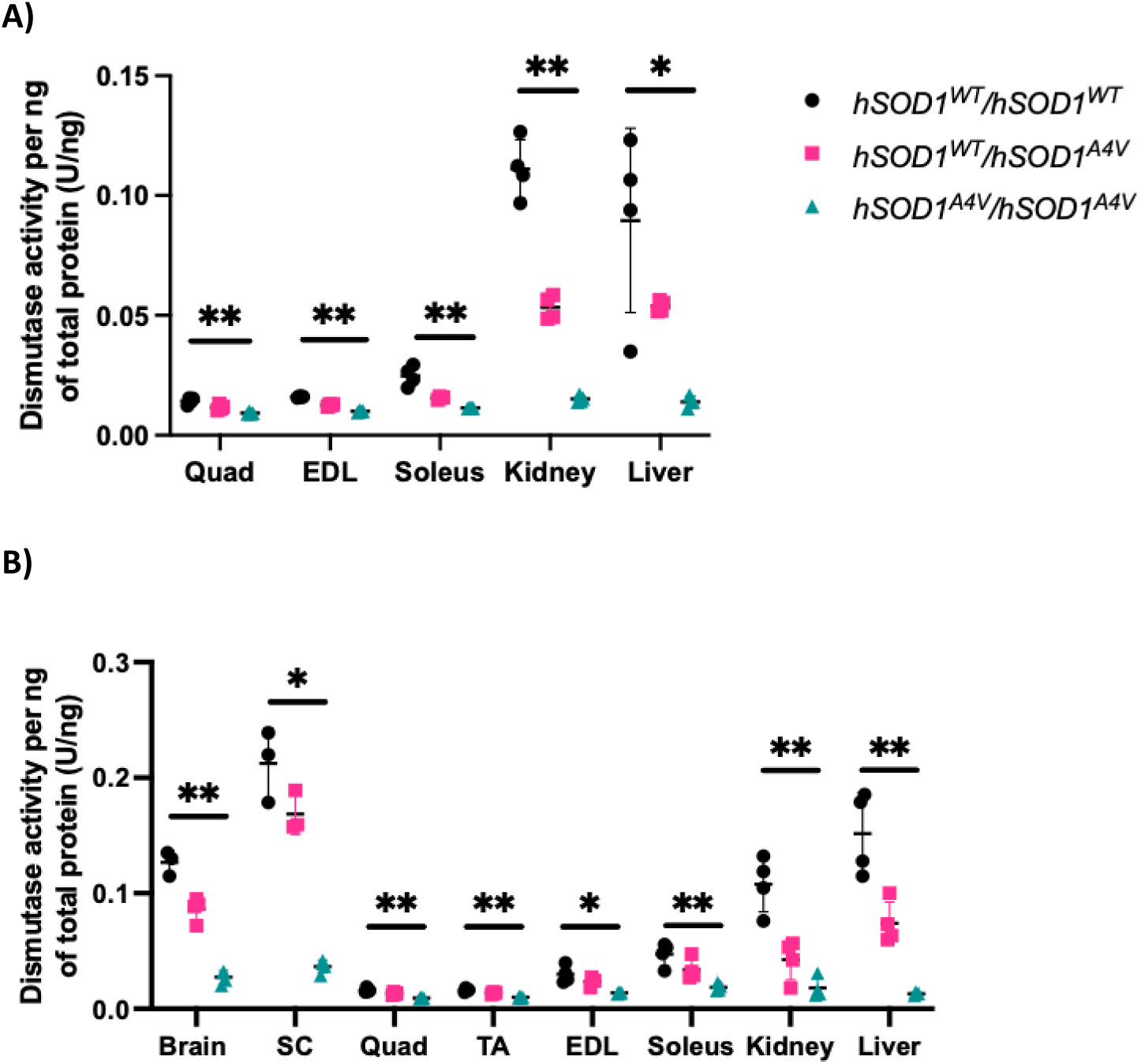
SOD1 activity in *hSOD1^A4V^* mice. A) Spectrophotometric dismutase activity assays in tissues from male 3-month-old *hSOD1^A4V^* mice. Mean ± SEM. One-way ANOVA with Tukey’s post hoc test. \**P* < 0.05, \*\**P* < 0.01, \*\*\**P* < 0.001, \*\*\*\**P* < 0.0001. B) Spectrophotometric dismutase activity assays in tissues from female 3-month-old *hSOD1^A4V^* mice. Mean ± SEM. One-way ANOVA with Tukey’s post hoc test. \**P* < 0.05, \*\**P* < 0.01, \*\*\**P* < 0.001, \*\*\*\**P* < 0.0001.

**Supplementary Figure 5.**
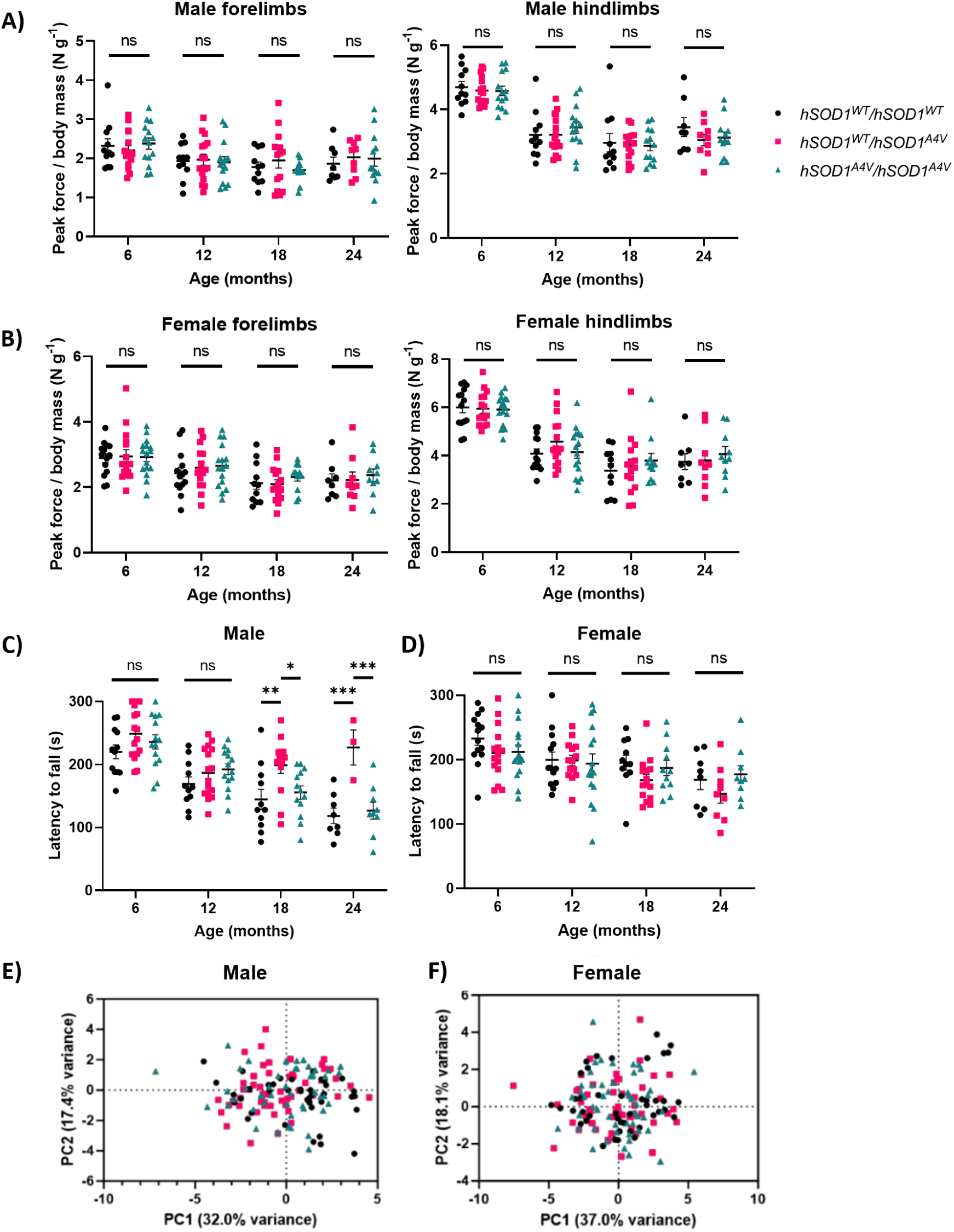
Motor function phenotyping in *hSOD1^A4V^* mice. A) Grip strength data from male forelimb and hindlimb (n=8-16 per genotype) *hSOD1^A4V^* mice. B) Grip strength data from female forelimb and hindlimb (n=9-16 per genotype) *hSOD1^A4V^*mice. C) Rotarod data from male (n=8-16 per genotype) *hSOD1^A4V^*mice. Mean ± SEM. Two-way ANOVA with Tukey’s post hoc test. Reported significance values are from the Tukey multiple comparisons test. *P < 0.05. **P < 0.01, ***P < 0.001. ns, not significant. D) Rotarod data from female (n=9-16 per genotype) *hSOD1^A4V^* mice. Mean ± SEM. Two-way ANOVA with Tukey’s post hoc test. Reported significance values are from the Tukey multiple comparisons test. *P < 0.05. **P < 0.01, ***P < 0.001. ns, not significant. E) Principal component analysis (PCA) using all 13 gait analysis parameters (walking speed, forelimb and hindlimb stance duration, step length, stride length, stride width, swing duration and duty factor) as variables in male hSOD1^A4V^ mice. F) Principal component analysis (PCA) using all 13 gait analysis parameters (walking speed, forelimb and hindlimb stance duration, step length, stride length, stride width, swing duration and duty factor) as variables in female *hSOD1^A4V^* mice.

**Supplementary Figure 6.**
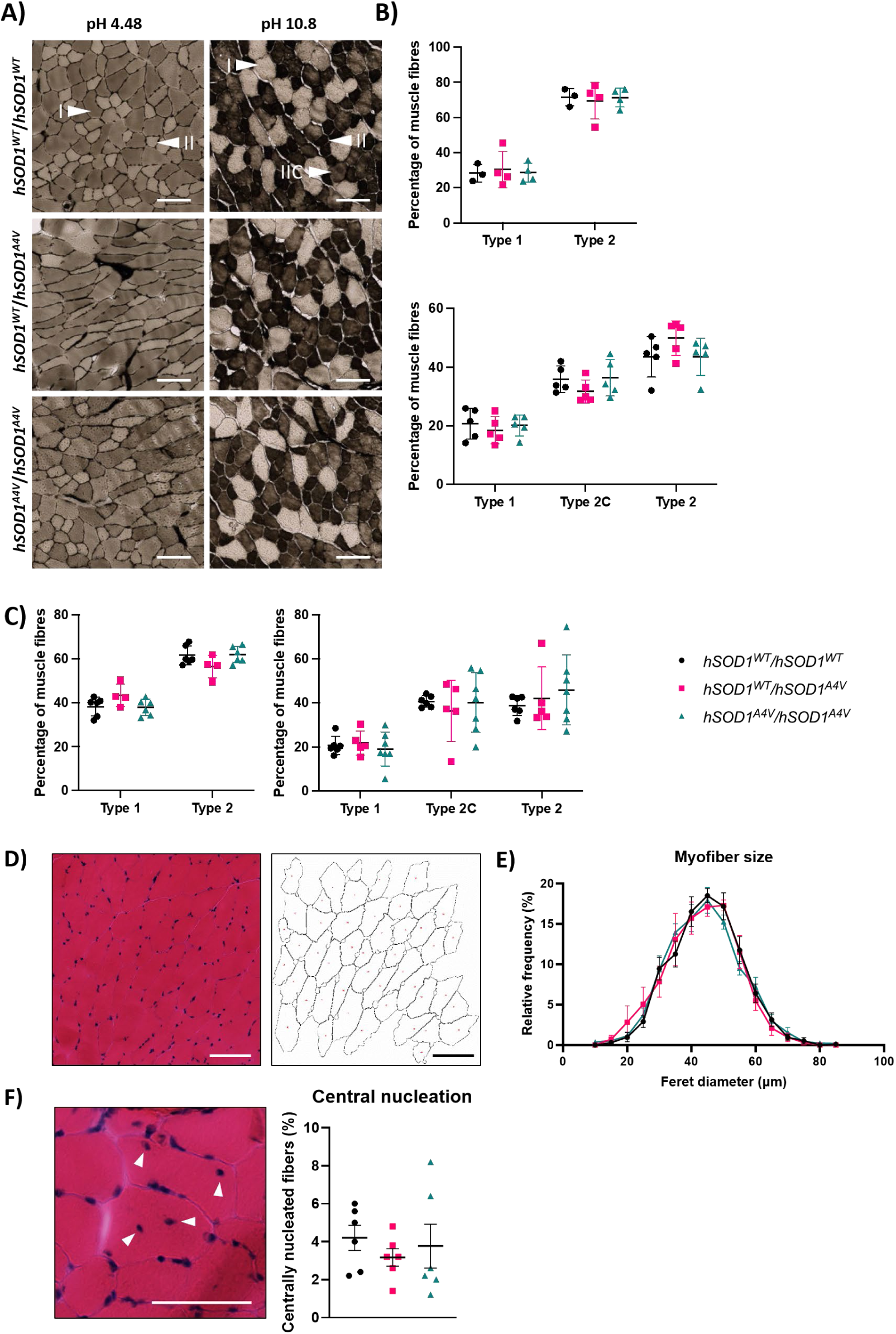
Myofibre typing and analysis in *hSOD1^A4V^* mice. A) Representative images of myosin-ATPase staining in TA muscle from 6-month-old female *hSOD1^A4V^* mice. Arrowheads indicate type I, type II and type IIC muscle fibres. Scale bar = 100μm. B) Quantification of myofiber type in TA muscle from 6-month-old female *hSOD1^A4V^*mice (n=5 per genotype). C) Quantification of myofiber type in TA muscle from 27-month-old female *hSOD1^A4V^* mice (n=5-7 per genotype). D) Representative image of a H&E-stained TA muscle section and the corresponding segmented image used for Feret diameter determination. Scale bar = 100μm. E) Quantification of myofiber size in TA muscle from 27-month-old female *hSOD1^A4V^* mice (n=5 per genotype), measured by the minimum Feret diameter. F) Example of centrally nucleated myofibers. Arrowheads indicate centralised nuclei. Scale bar = 50μm. Percentage of centrally nucleated myofibers in TA muscle from 27-month-old female *hSOD1^A4V^*mice (n=6 per genotype).

**Supplementary Figure 7.**
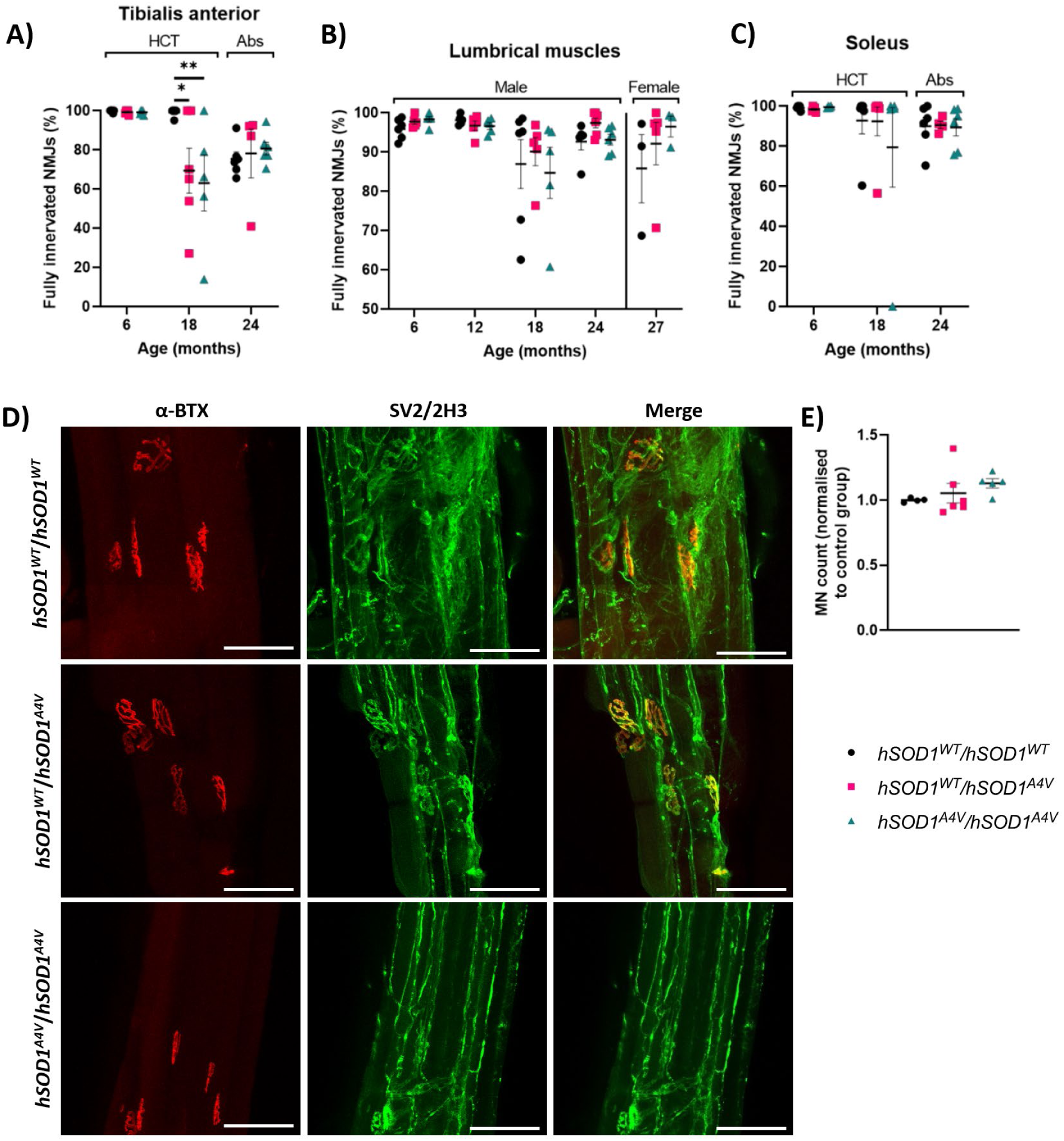
Neuromuscular junction analysis and motor neuron counts in *hSOD1^A4V^* mice. A) Quantification of fully innervated NMJs in lumbrical muscles from 24-month-old male *hSOD1^A4V^* mice (n=4-6 animals per genotype). B) Quantification of fully innervated NMJs in tibialis anterior muscles from 24-month-old male (n=5-6 animals per genotype) and female (n=3-5 animals per genotype) *hSOD1^A4V^* mice. The axons and presynaptic termini were visualised using antibodies against synaptophysin/neurofilament. C) Quantification of fully innervated NMJs in soleus muscle from 24-month-old male *hSOD1^A4V^* mice (n=4-6 animals per genotype). Two-way ANOVA followed by the Tukey multiple comparisons test. Reported significance values are from Tukey’s test. Mean ± SEM. \**P* < 0.05. **P < 0.01. Abs, antibodies. D) Representative images of neuromuscular junction (NMJ) staining in lumbrical muscles from 24-month-old *hSOD1A4V* mice. α-bungarotoxin (α-BTX), red; synaptophysin/neurofilament (SV2/2H3), green. Scale bar 100 μm. E) Quantification of motor neurons within the sciatic pool of the lumbar spinal cord (L3-L5) of 24-month-old male *hSOD1^A4V^* animals. n=4-6 animals per genotype. Mean ± SEM.

**Supplementary Figure 8.**
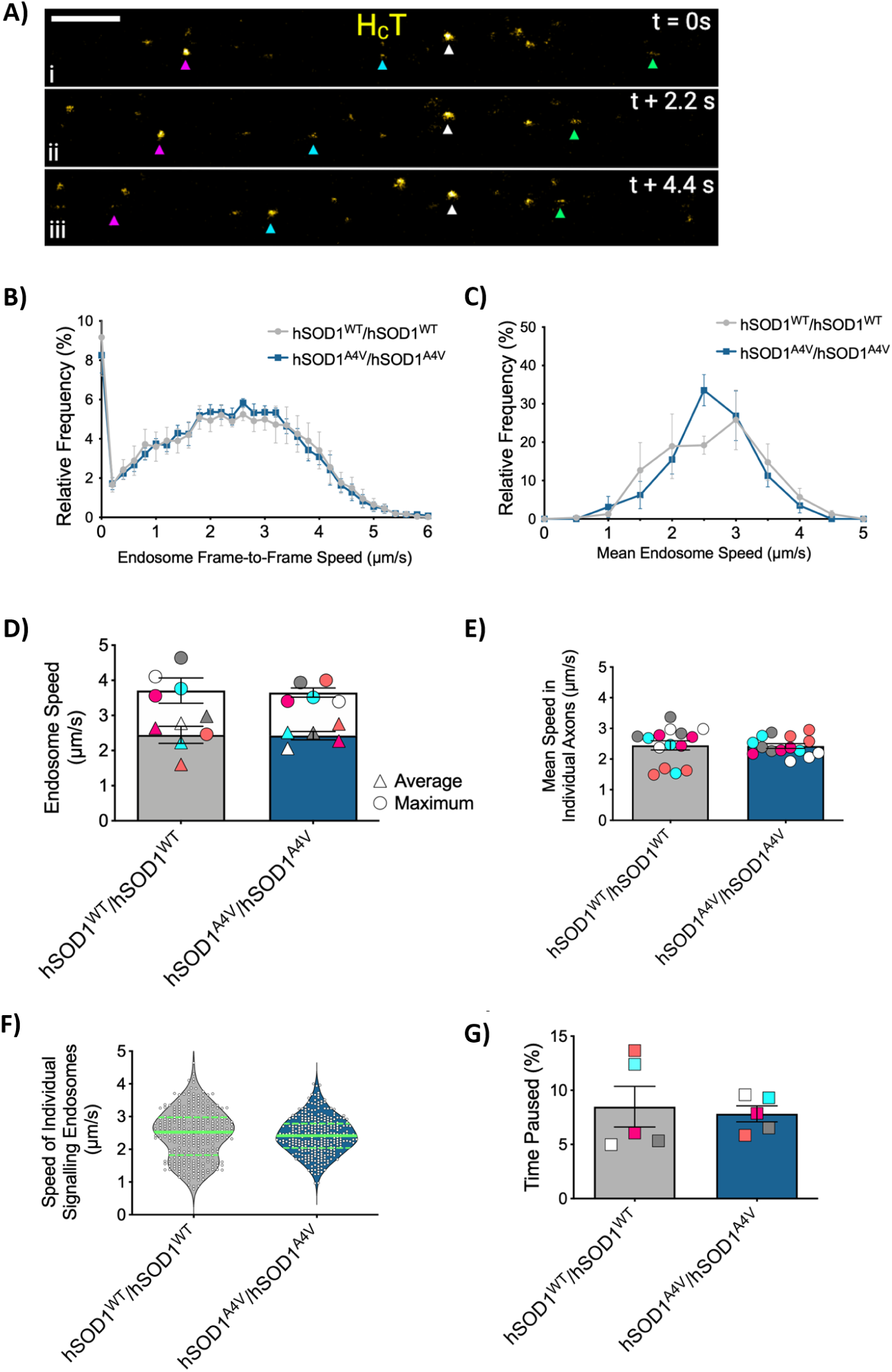
Axonal trafficking analysis in *hSOD1^A4V^*mice. A) Retrogradely transported H_C_T-positive endosomes (pseudo-coloured yellow) from tibialis anterior innervating motor axons are individually tracked to assess their dynamics. Colour-coded arrowheads (cyan, green and magenta) identify three individual endosomes that are undergoing retrograde axonal transport, whereas the white arrowheads identify a stationary/paused endosome. Scale bar = 5µm; t = 2.2 secs. *hSOD1^WT^*and *hSOD1^A4V^* mice displayed no differences in *in vivo* axonal transport dynamics as demonstrated by the B) endosomal frame-to-frame; and C) mean endosome distribution curves; D) average signalling endosome speeds per animal (maximum [circles] P = 0.55, mean [triangles] P = 0.69, Mann Whitney); E) mean speeds in individual axons (P = 0.49, Mann Whitney), F) individual signalling endosome speeds (green lines indicate the median and the green dashed lines represent the upper and lower quartiles); and G) pausing (P = 0.84, Mann Whitney).

**Supplementary Figure 9.**
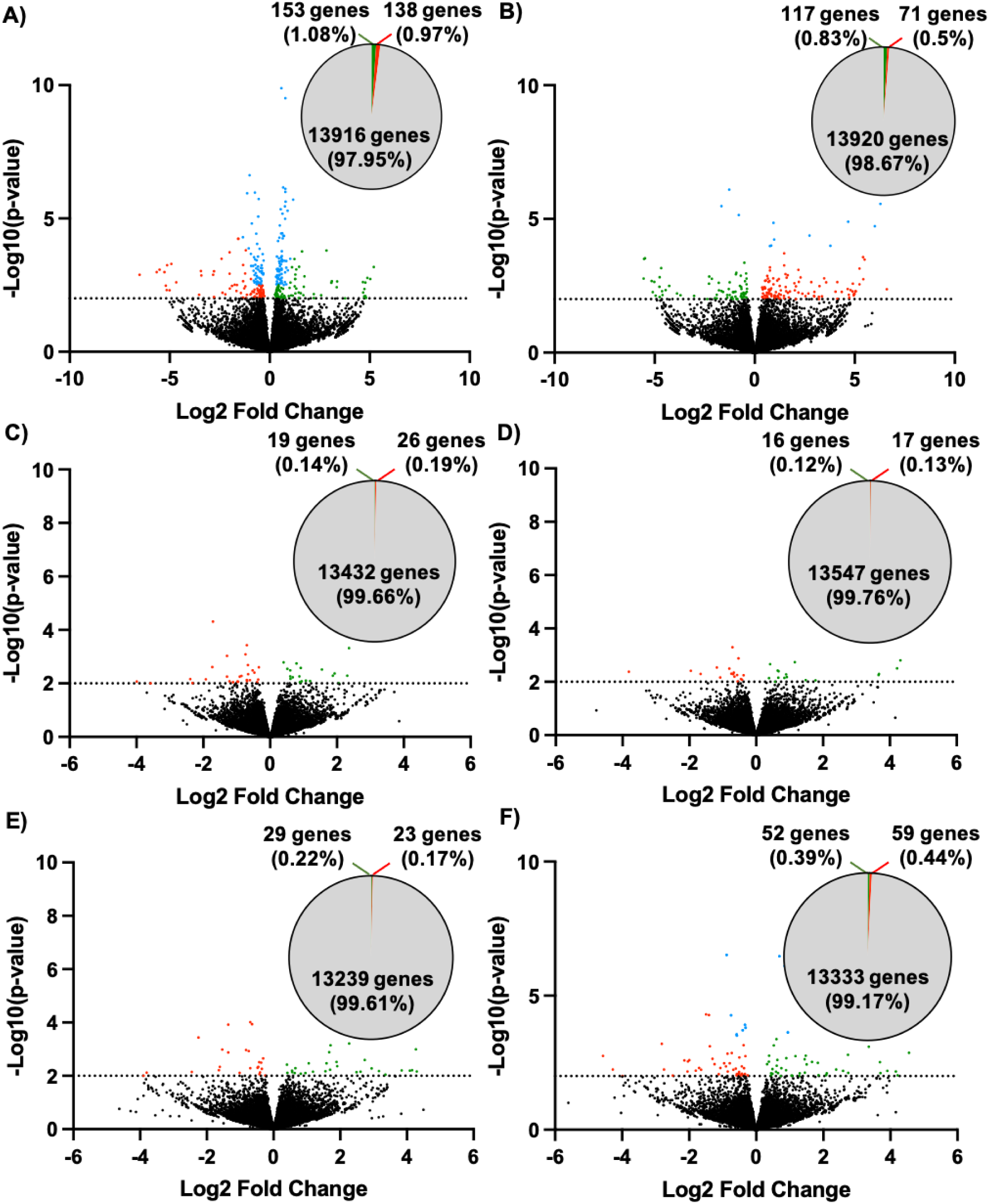
Differential gene expression in the tibialis anterior of *hSOD1^A4V^*mice compared to *hSOD1^WT^* controls. Volcano plots showing significantly upregulated genes (green) and downregulated genes (red) (p value < 0.01, log2fold change > 1.3 or <-1.3) in the tibialis anterior of 6-month-old A) *hSOD1^WT/A4V^* and B) *hSOD1^A4V/A4V^*, 12-month-old C) *hSOD1^WT/A4V^* and D) *hSOD1^A4V/A4V^* and 20-month-old E) *hSOD1^WT/A4V^* and F) male *hSOD1^A4V/A4V^* animals. Genes with p value < 0.01 but a log2fold change < 1.3 or >.1.3 are shown in blue. Non-significant genes with a p value > 0.01 are shown in black. Venn diagrams display the number of identified genes and the proportion that are significantly up-and down-regulated.

**Supplementary Figure 10.**
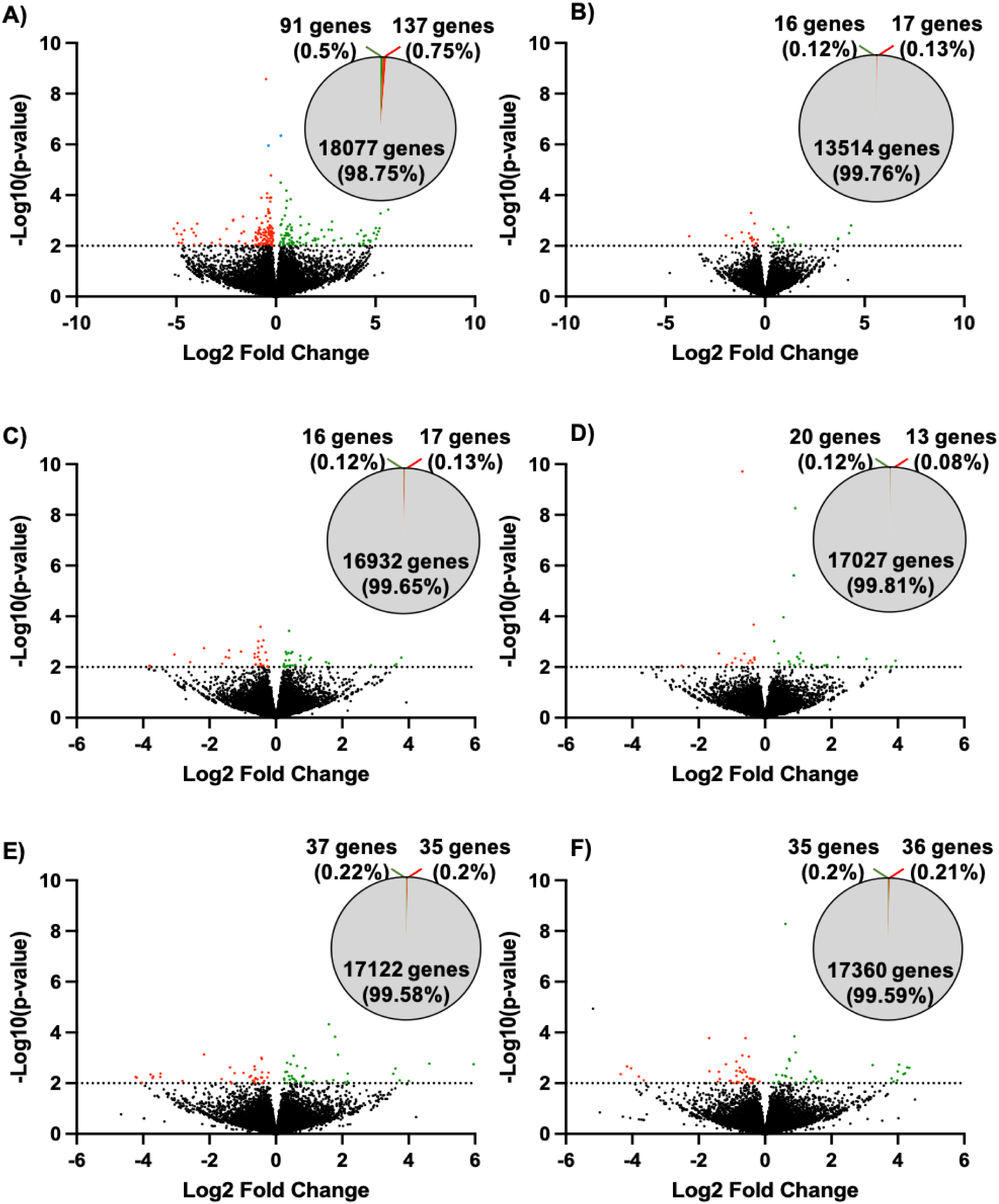
Differential gene expression in the spinal cord of *hSOD1^A4V^* mice compared to *hSOD1^WT^* controls. Volcano plots showing significantly upregulated genes (green) and downregulated genes (red) (p value < 0.01, log2fold change > 1.3 or <-1.3) in the spinal cord of 6-month-old A) *hSOD1^WT/A4V^* and B) *hSOD1^A4V/A4V^*, 12-month-old C) *hSOD1^WT/A4V^* and D) *hSOD1^A4V/A4V^* and 20-month-old E) *hSOD1^WT/A4V^* and F) male *hSOD1^A4V/A4V^*animals. Genes with p value < 0.01 but a log2fold change < 1.3 or >.1.3 are shown in blue. Non-significant genes with a p value > 0.01 are shown in black. Venn diagrams display the number of identified gens and the proportion that are significantly up and down regulated.

**Supplementary Figure 11.**
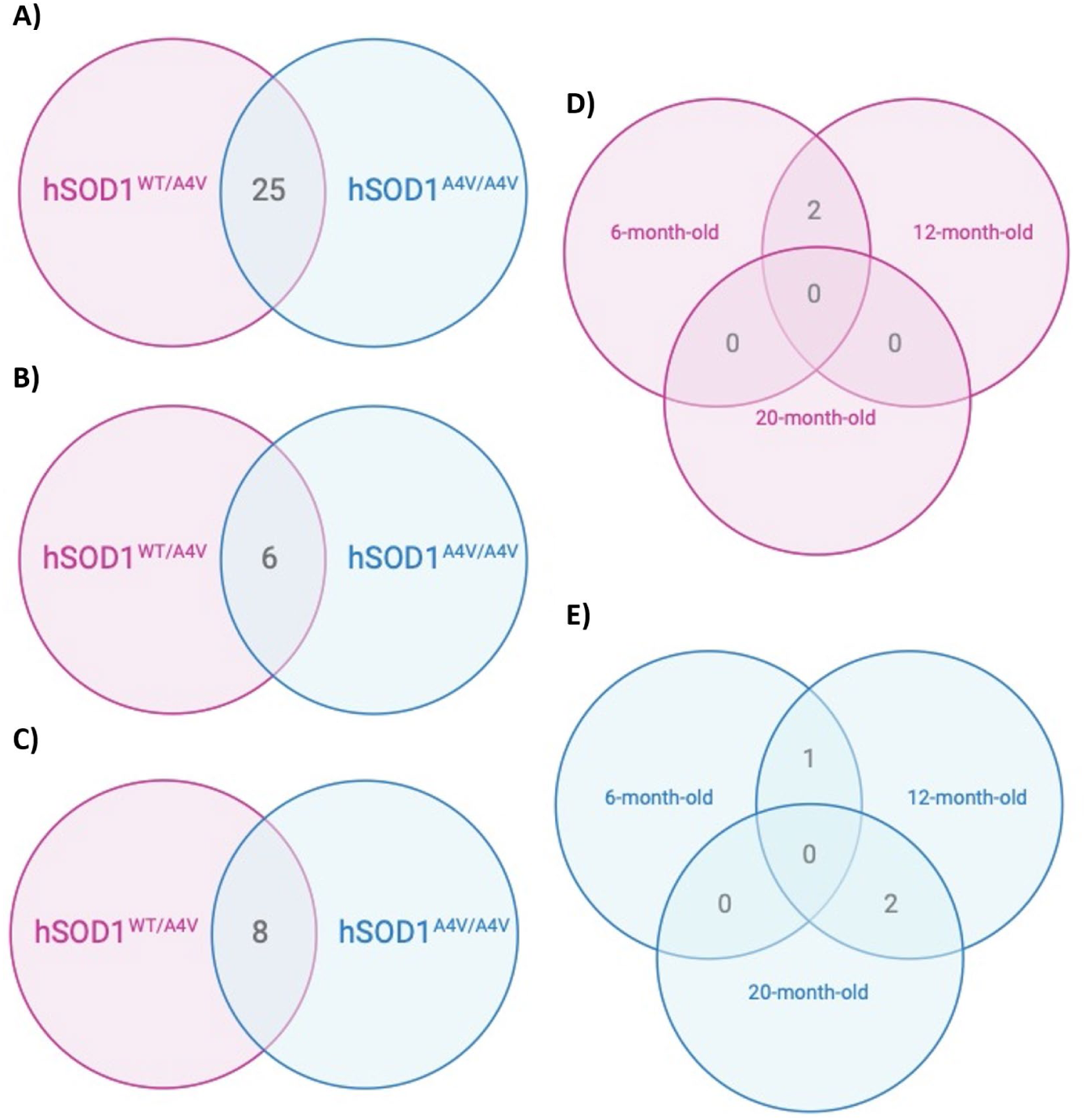
Overlap of differentially expressed genes the tibialis anterior of *hSOD1^A4V^*mice compared to *hSOD1^WT^* controls. Venn plots displaying the overlap of common differentially expressed genes in male *hSOD1^WT/A4V^* (pink) and *hSOD1^A4V/A4V^* (blue) animals in A) 6-month-old B) 12-month-old and C) 20-month-old animals. D) Common differentially expressed genes at 6-, 12-and 20-months-old in *hSOD1^WT/A4V^* animals. E) Common differentially expressed genes at 6-, 12-and 20-months-old in *hSOD1^A4V/A4V^*animals.

**Supplementary Figure 12.**
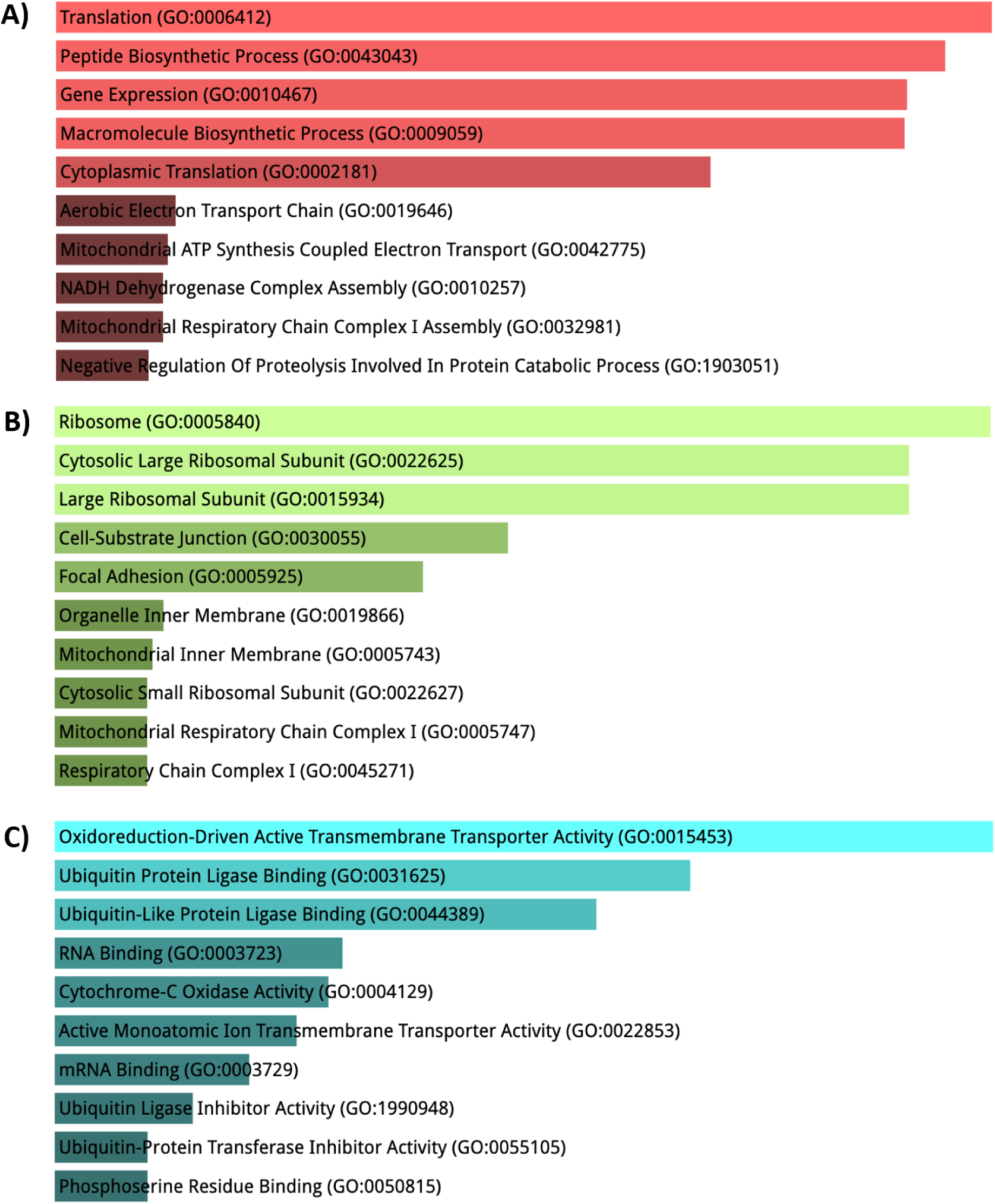
Enriched GO terms associated with significantly differentially expressed genes in the TA of *hSOD1^WT/A4V^*mice. Analysis using combined up-and downregulated datasets from differentially expression analysis showed biological process (A), cellular components (B) and molecular functions (C) most affected in male of 6-, 12-and 20-month old *hSOD1^WT/A4V^* mice.

**Supplementary Figure 13.**
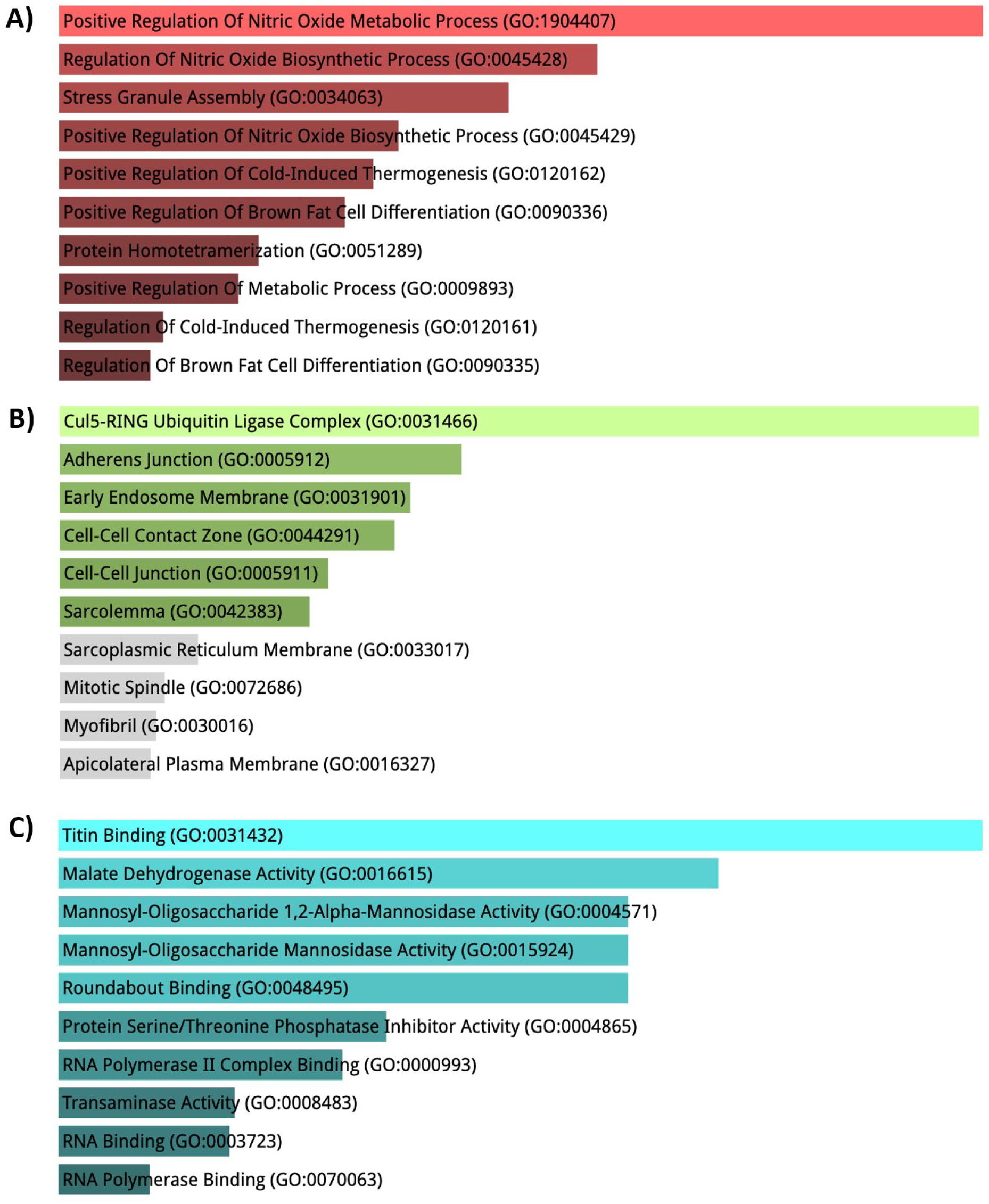
Enriched GO terms associated with significantly differentially expressed genes in the TA of *hSOD1^A4V/A4V^*mice. Analysis using combined up-and downregulated datasets from differentially expression analysis showed biological process (A), cellular components (B) and molecular functions (C) most affected in of 6-, 12-and 20-month old male *hSOD1^A4V/A4V^* mice.

**Supplementary Figure 14.**
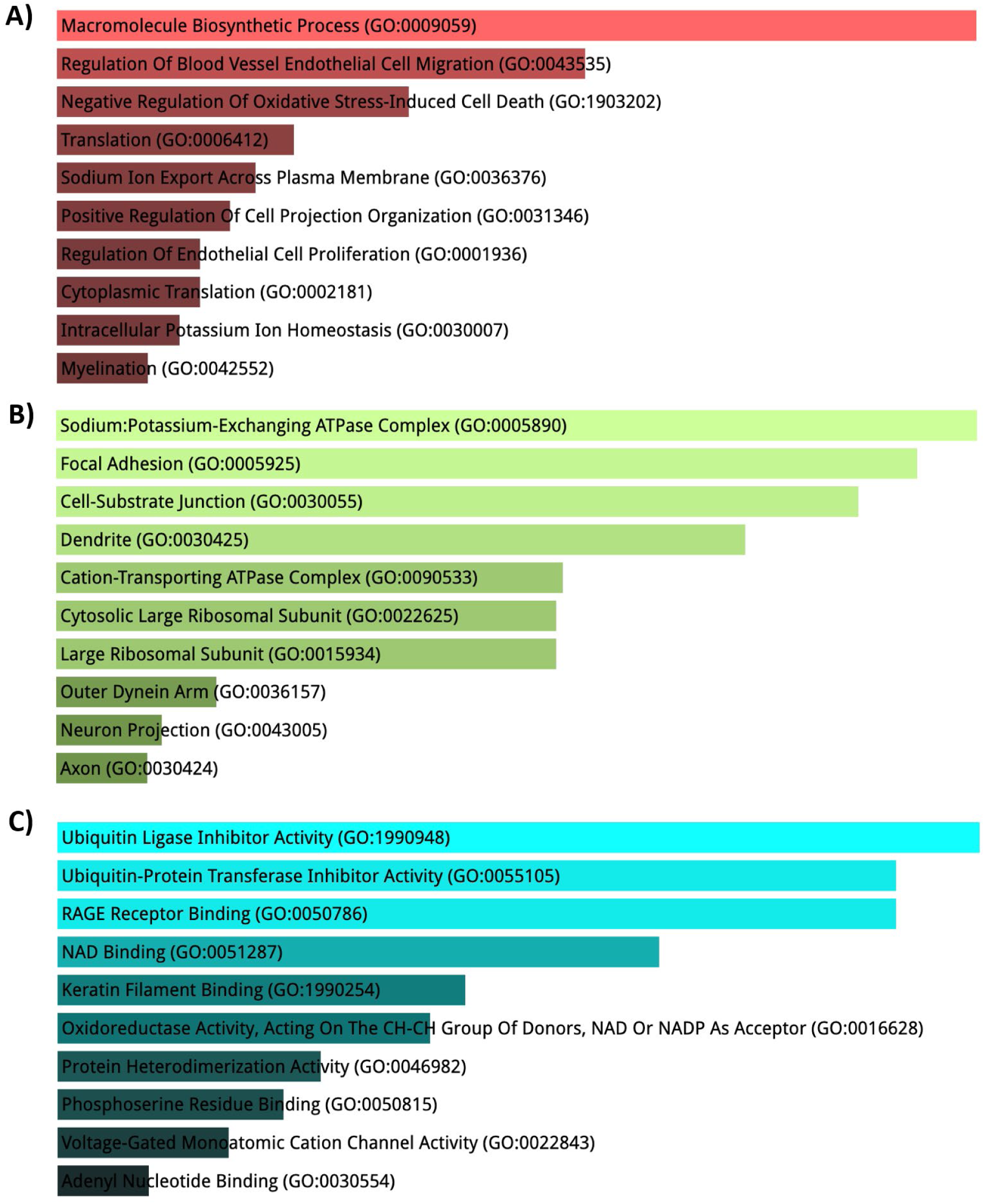
Enriched GO terms associated with significantly differentially expressed genes in the SC of *hSOD1^WT/A4V^*mice. Analysis using combined up-and downregulated datasets from differentially expression analysis showed biological process (A), cellular components (B) and molecular functions (C) most affected in of 6-, 12-and 20-month old male *hSOD1^WT/A4V^* mice.

**Supplementary Figure 15.**
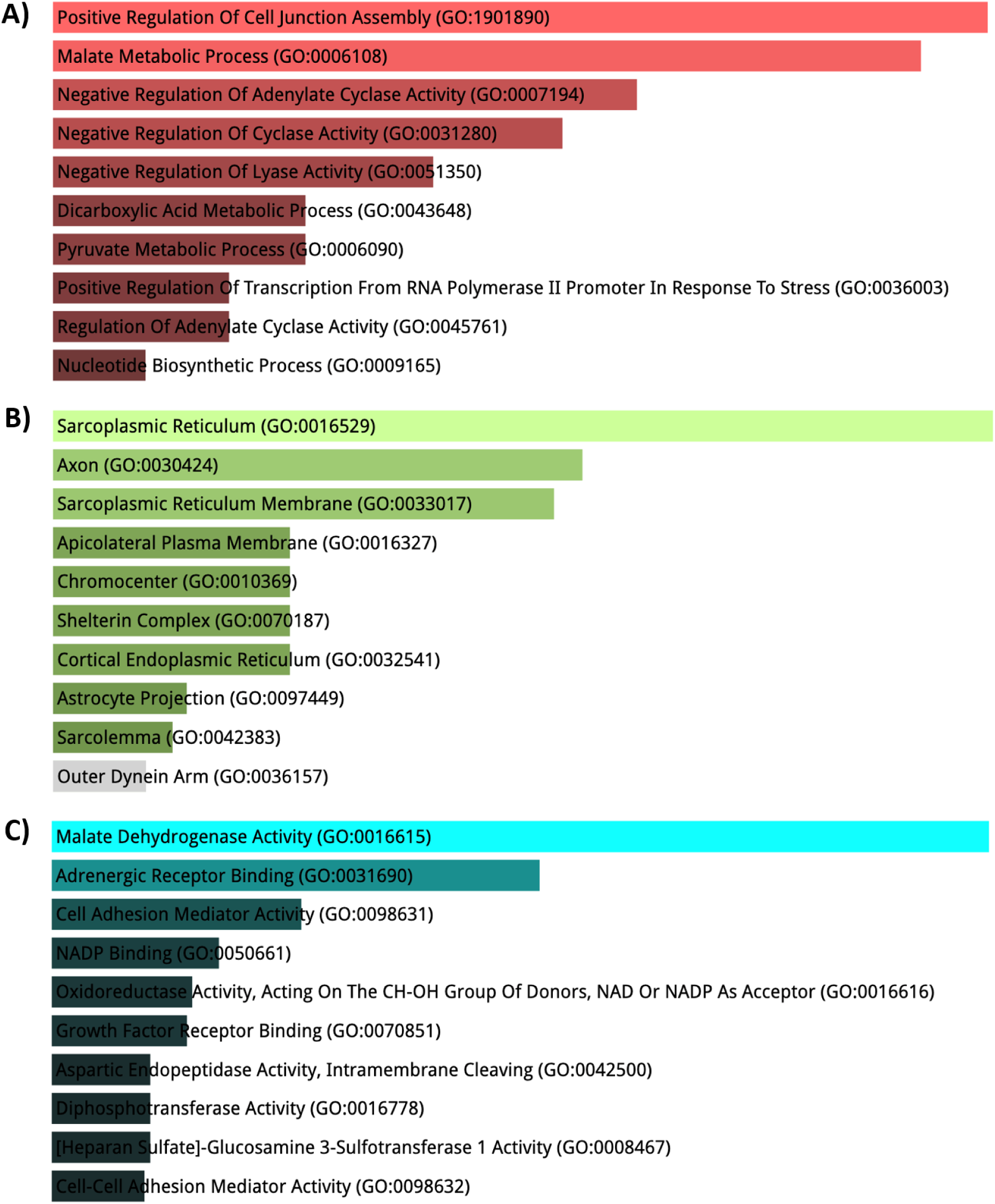
Enriched GO terms associated with significantly differentially expressed genes in the SC of *hSOD1^A4V/A4V^*mice. Analysis using combined up-and downregulated datasets from differentially expression analysis showed biological process (A), cellular components (B) and molecular functions (C) most affected in of 6-, 12-and 20-month old male *hSOD1^A4V/A4V^* mice.

**Supplementary Figure 16.**
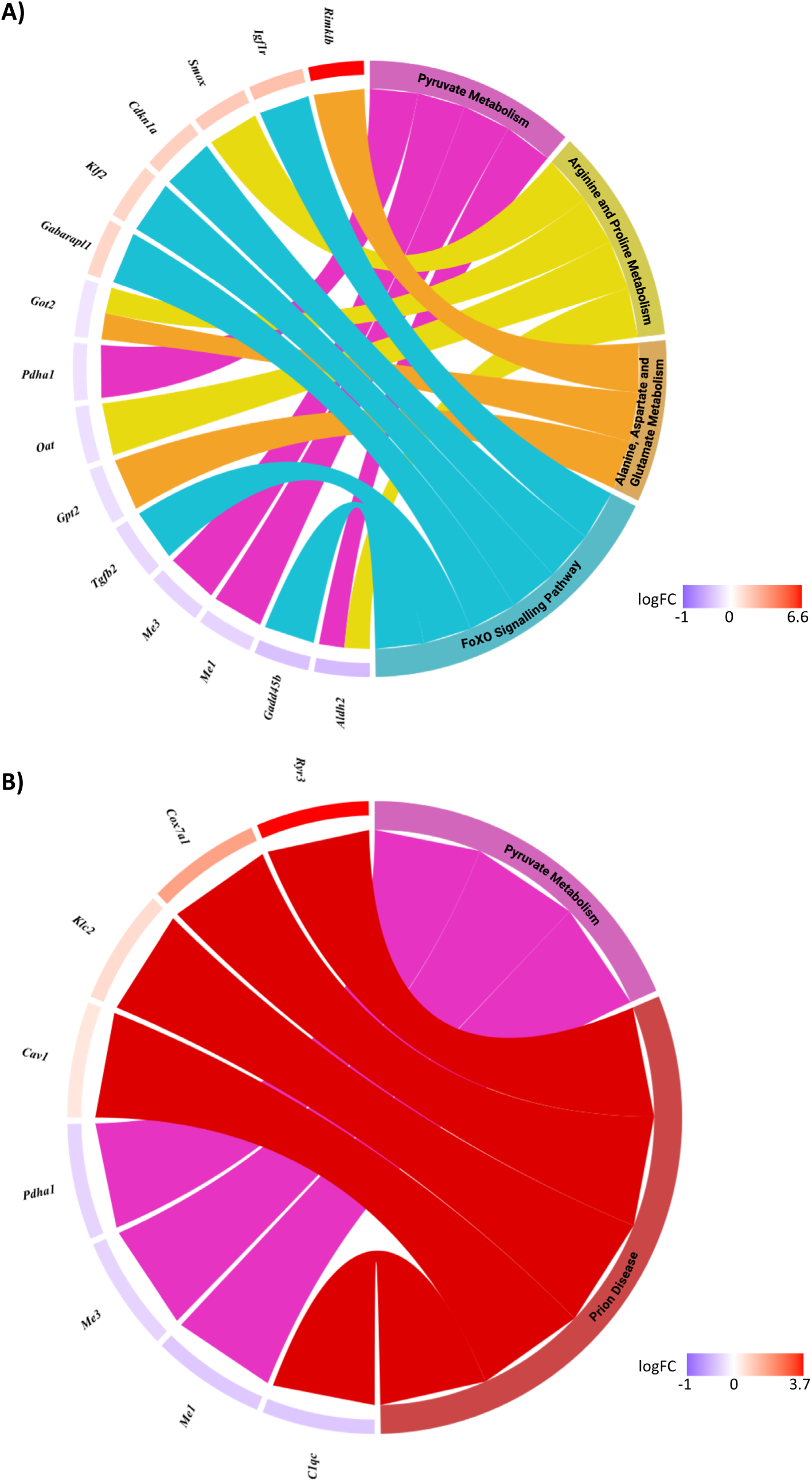
Chord diagram showing the core genes of significant affected pathways in *hSOD1^WT/A4V^* mice compared to littermate controls. Significant pathways identified from DEGs in the A) TA and B) SC of *hSOD1^WT/A4V^* vs *hSOD1^WT/WT^* of 6-, 12-and 20-month old male mice. Significant pathways are shown on the right and the fold change of core genes is shown on the left. Left-right connections indicate gene membership in a pathways leading-edge subset. Image constructed using Biorender.

**Supplementary Figure 17.**
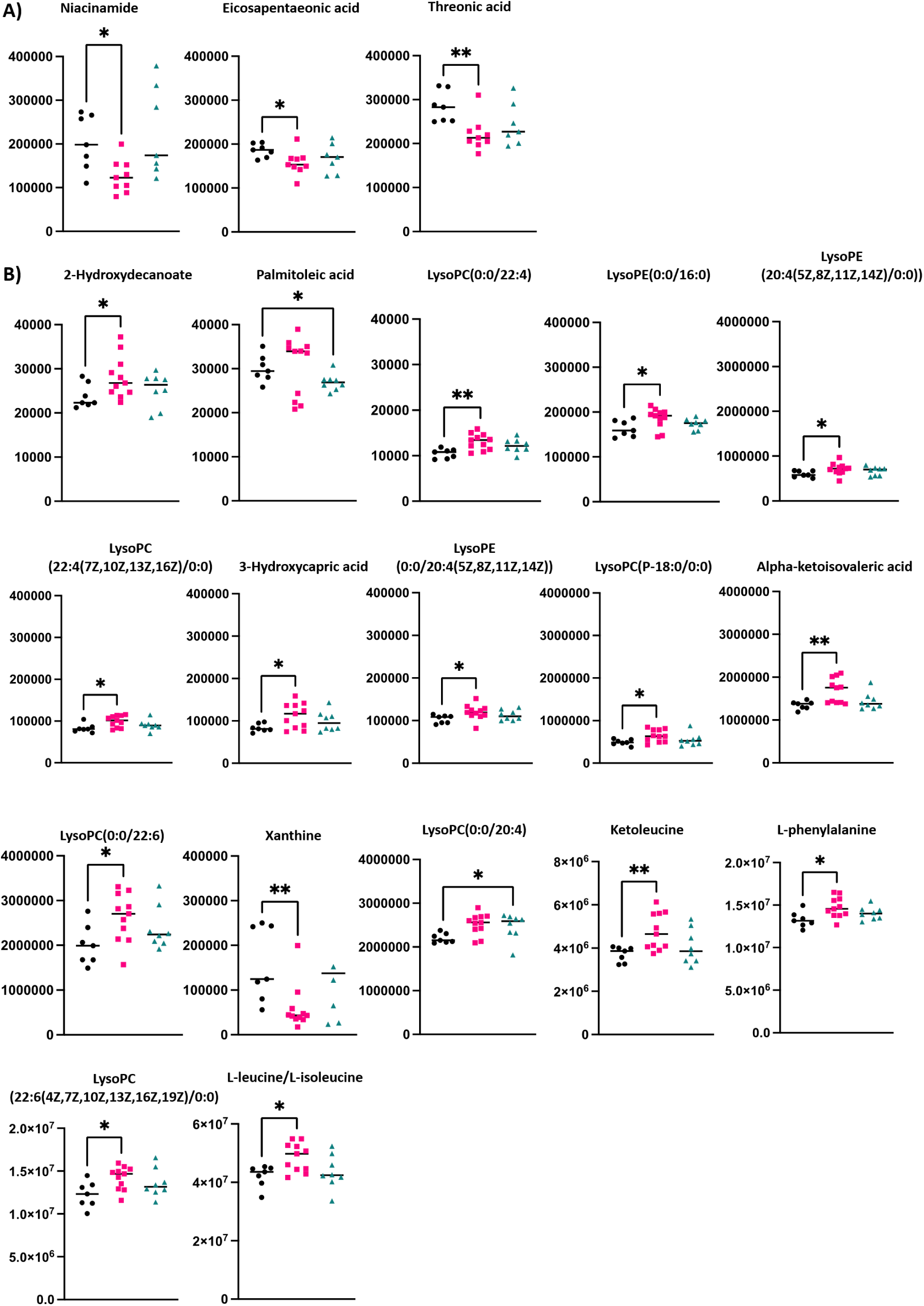
Metabolic changes in plasma of 3-and 6-month-old *hSOD1^A4V^*mice. Significantly altered metabolites in the blood of male and female A) 3- and B) 6- month-old *hSOD1^WT/A4V^* and *hSOD1^A4V/A4V^*animals compared to littermate controls. One-way ANOVA with Kruskal-Wallis test for multiple comparisons was used to determine significance with *pvalue < 0.05, ** pvalue < 0.01.

**Supplementary Figure 18.**
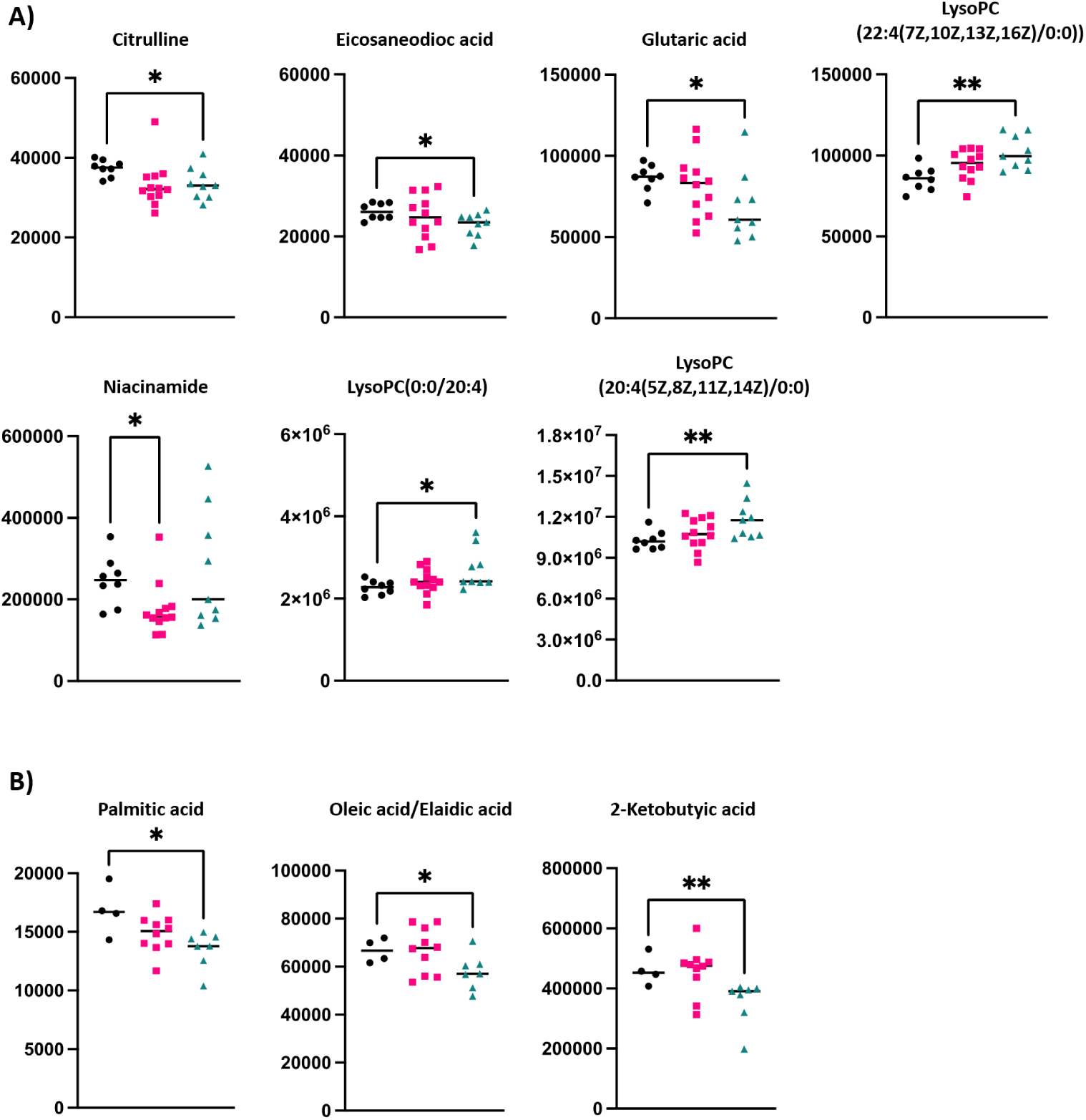
Metabolic changes in plasma of 9 and 12-month-old *hSOD1^A4V^* mice. Significantly altered metabolites in the blood of male and female A) 9- and B) 12-month-old *hSOD1^WT/A4V^* and *hSOD1^A4V/A4V^*animals compared to littermate controls. One-way ANOVA with Kruskal-Wallis test for multiple comparisons was used to determine significance with *p value < 0.05, ** p value < 0.01.

**Supplementary Figure 19.**
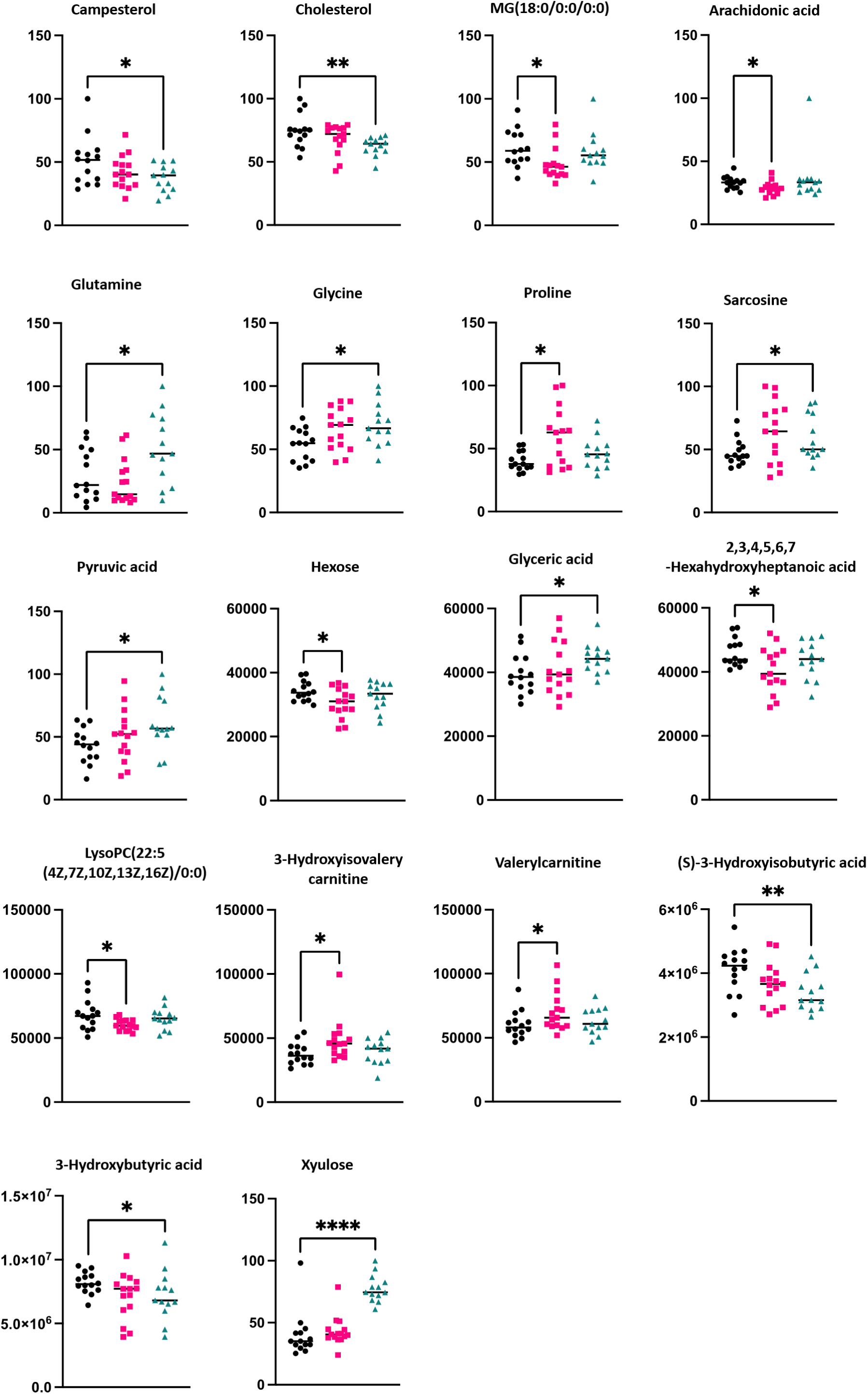
Metabolic changes in plasma of 20-month-old *hSOD1^A4V^* mice. Significantly altered metabolites in the blood of male and female 20-month-old *hSOD1^WT/A4V^*and *hSOD1^A4V/A4V^* animals compared to littermate controls. One-way ANOVA with Kruskal-Wallis test for multiple comparisons was used to determine significance with *pvalue < 0.05, ** pvalue < 0.01, **** pvalue < 0.001.

**Supplementary Figure 20.**
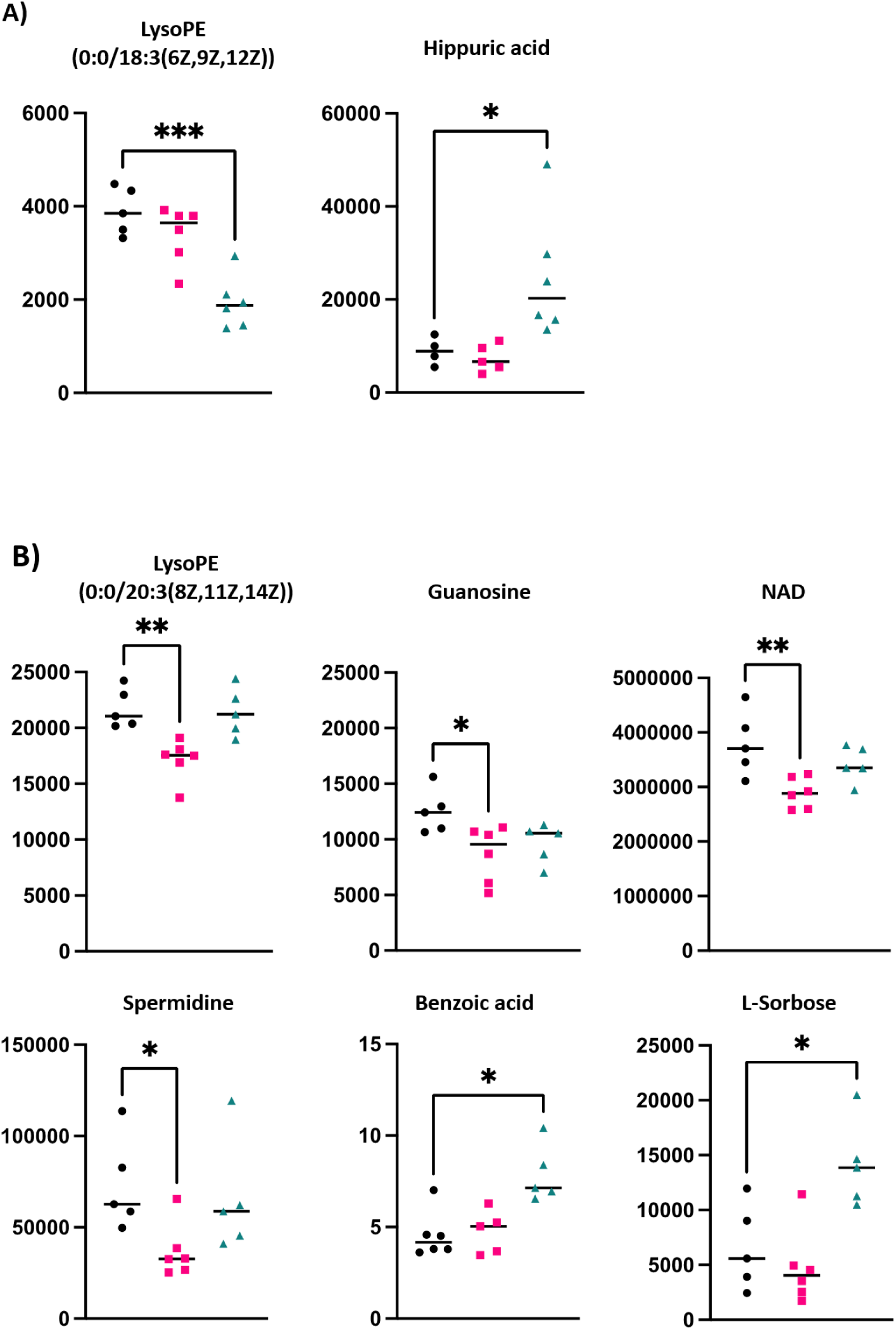
Metabolic changes in the spinal cord and tibialis anterior of 20-month-old *hSOD1^A4V^* mice. Significantly altered metabolites in the A) spinal cord and B) tibialis anterior of male 20-month-old *hSOD1^WT/A4V^* and *hSOD1^A4V/A4V^*animals compared to littermate controls. One-way ANOVA with Kruskal-Wallis test for multiple comparisons was used to determine significance with *p value < 0.05, ** p value < 0.01, *** p value < 0.005.

**Supplementary Figure 21.**
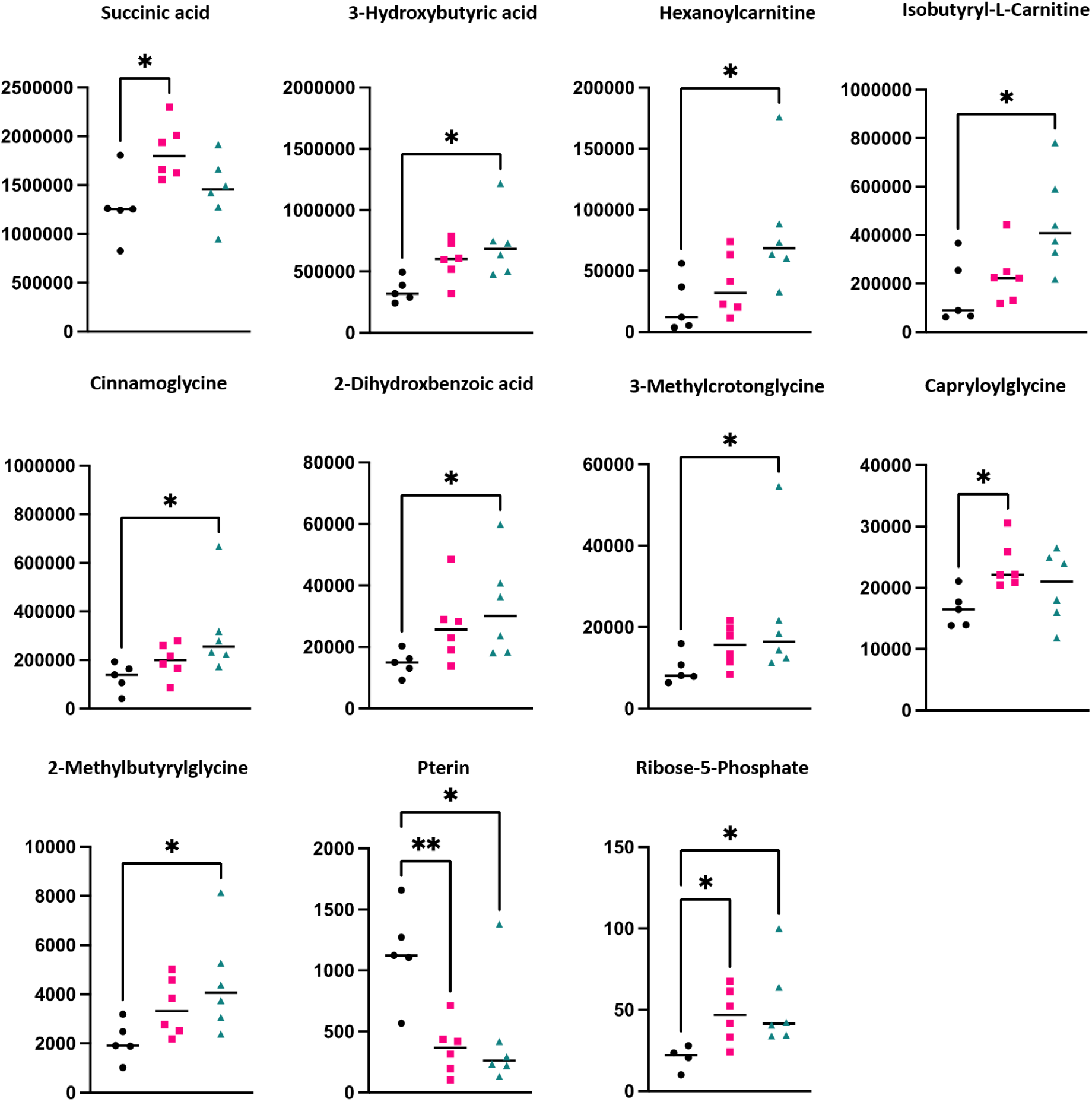
Metabolic changes in the liver of 20-month-old *hSOD1^A4V^* mice. Significantly altered metabolites in the liver of male 20-month-old *hSOD1^WT/A4V^*and *hSOD1^A4V/A4V^* animals compared to littermate controls. One-way ANOVA with Kruskal-Wallis test for multiple comparisons was used to determine significance with *p value < 0.05, ** p value < 0.01.

**Supplementary Figure 22.**
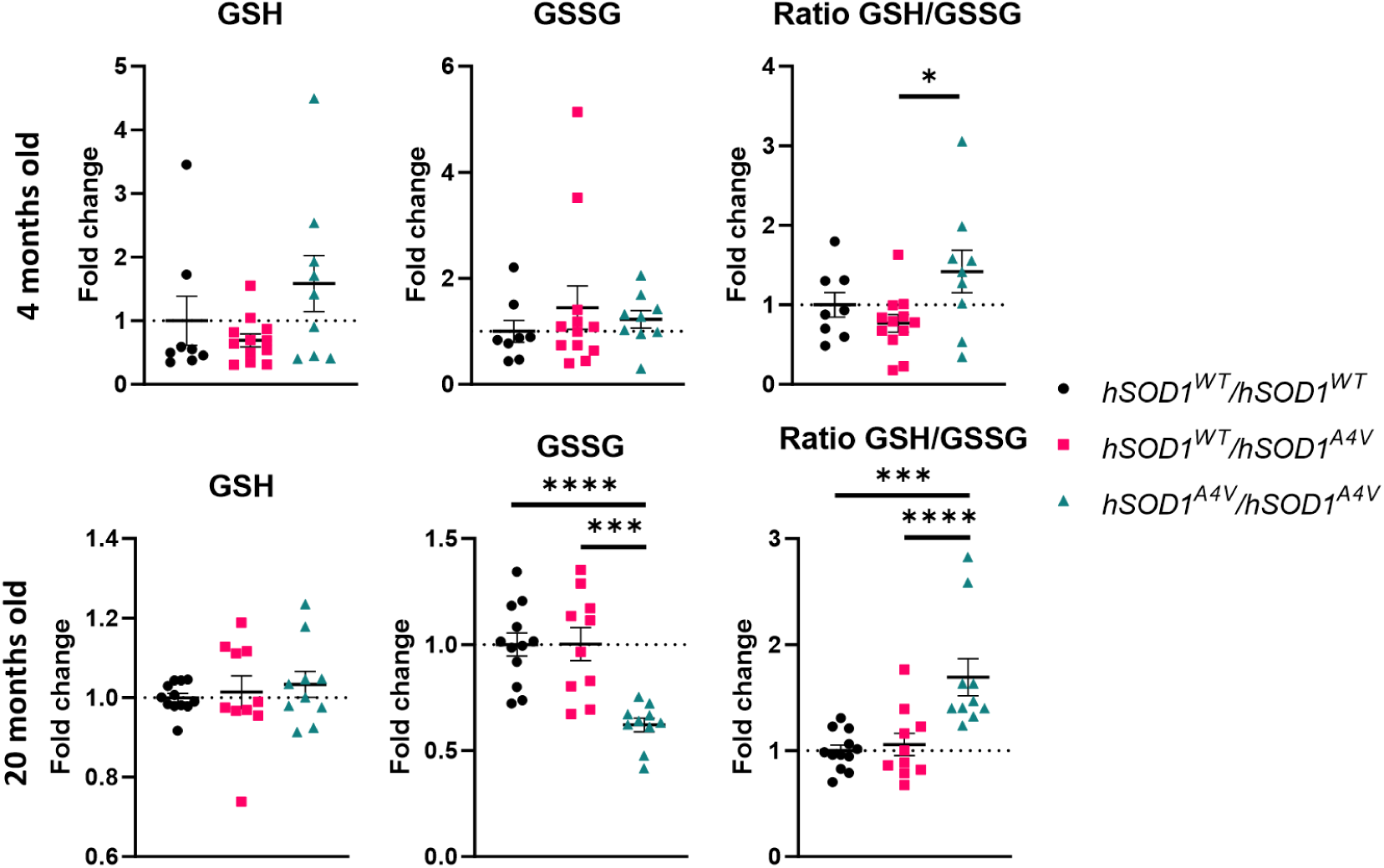
Blood glutathione measurements in *hSOD1^A4V^* mice. Blood glutathione measurements in 4- month-old and 20-month-old. Data are displayed as fold change relative to *hSOD1^WT/WT^* animals. 20-month-old data was pooled from two cohorts where each animal is displayed as fold change relative to *hSOD1^WT/WT^* animals from the same cohort. One-way ANOVA with Kruskal-Wallis test for multiple comparisons was used to determine significance with *p value < 0.05, ** p value < 0.01, *** p value < 0.005, **** p value < 0.001.

**Supplementary Figure 23.**
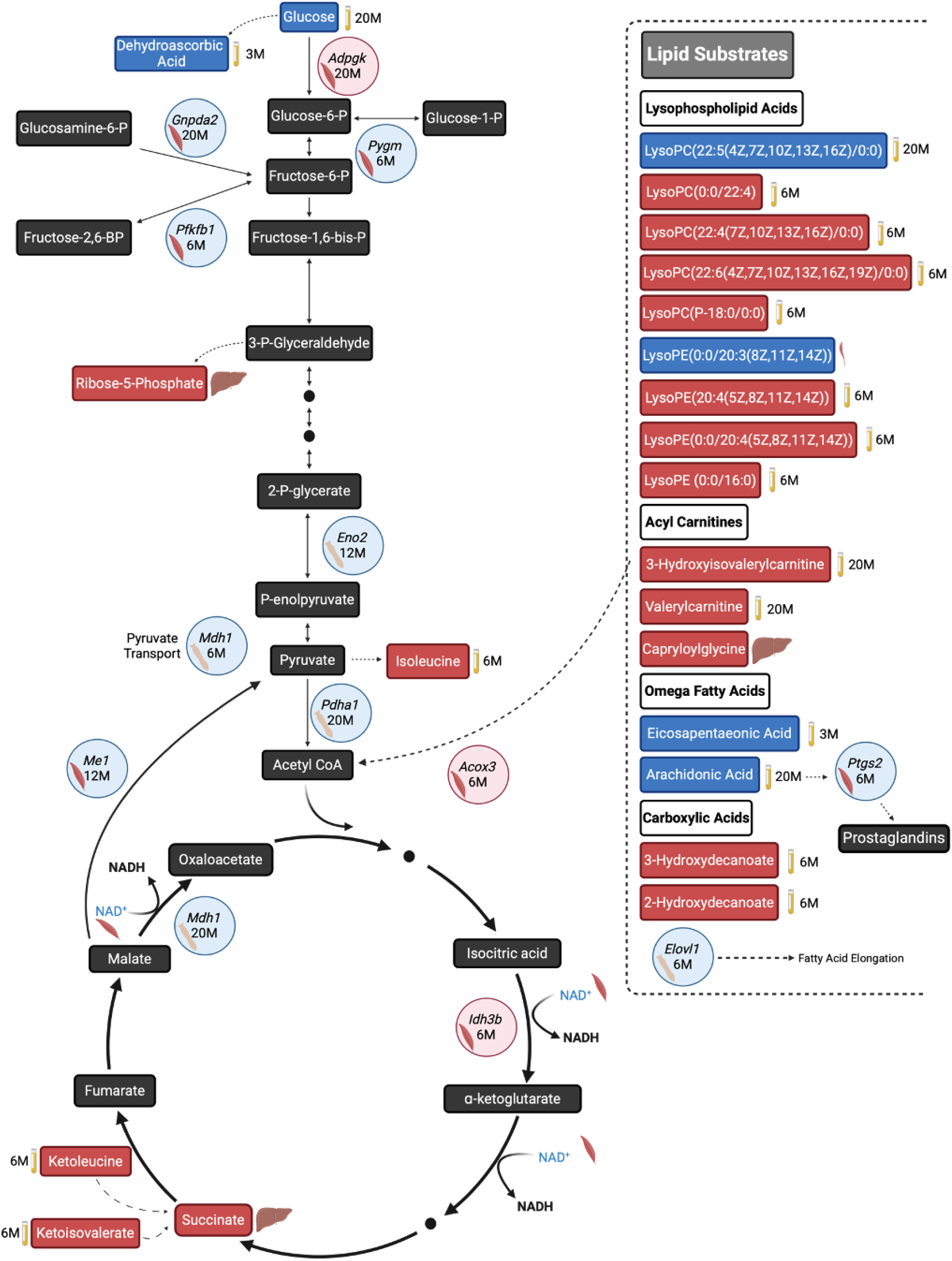
Integrated transcriptomic and metabolomic analysis of *hSOD1^WT/A4V^* mice highlights core energy metabolism pathways. Schematic representation of dysregulated energy metabolism pathways. Metabolites are depicted by rectangles, red: increased, blue: decreased and black: undetected. Genes are depicted by circles, red: increased, blue: decreased. Solid arrows indicate direct reaction, dotted arrow indicates the reaction is broken by one or more sub-reactions. The tissue in which genes or metabolites were significantly altered are depicted by the presence of a muscle, spinal cord or liver image. The timepoint at which a gene or metabolite was detected as significantly changed is indicated in months (M). Image constructed using Biorender.

**Supplementary Figure 24.**
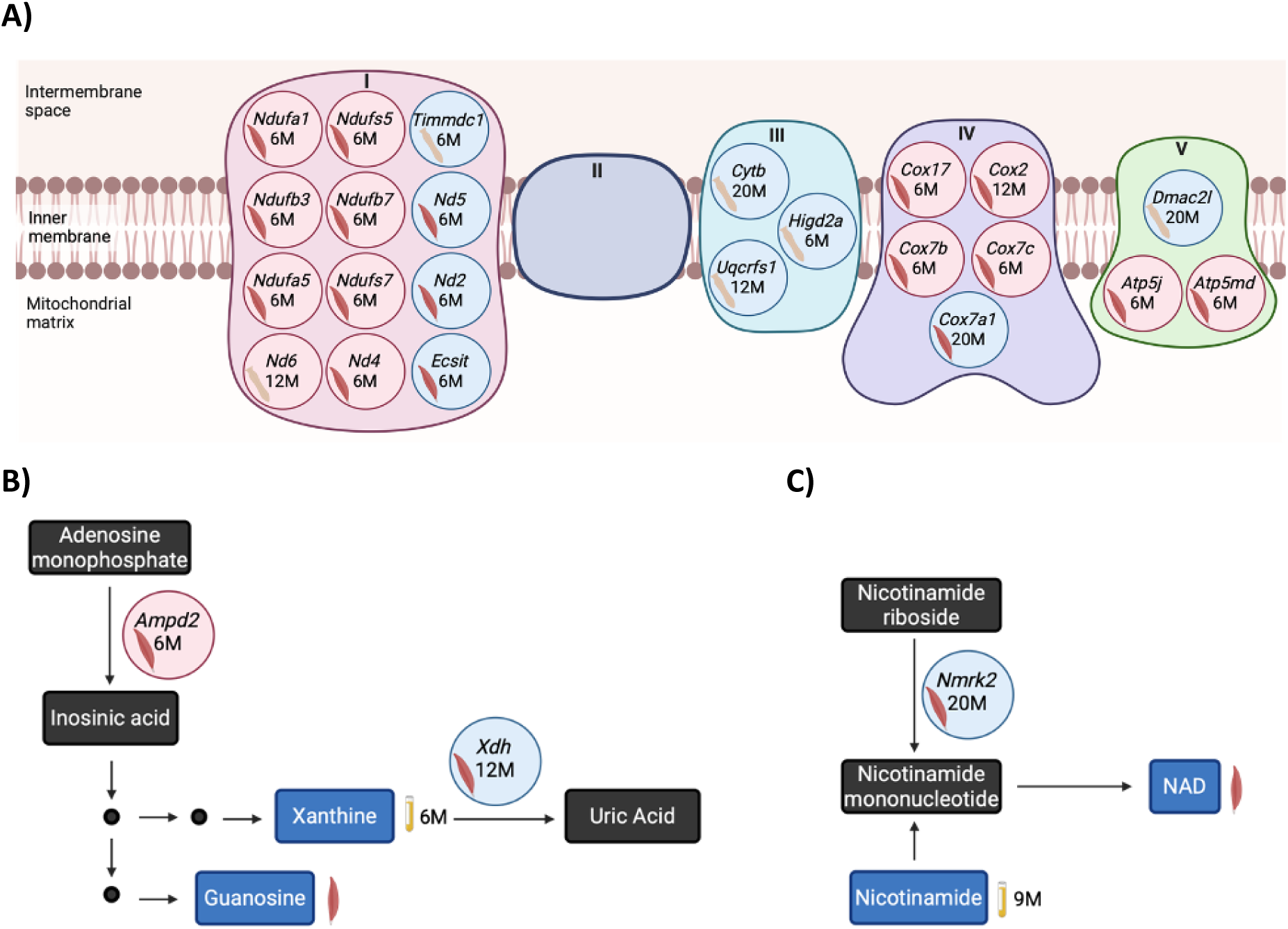
Integrated transcriptomic and metabolomic analysis of *hSOD1^WT/A4V^* mice highlights mitochondrial, xanthine and nicotinamide metabolic dysregulation. Schematic representation of dysregulated pathways, A) mitochondrial function, B) xanthine and C) NAD metabolism, in *hSOD1^WT/A4V^* mice. Metabolites are depicted by rectangles, red: increased, blue: decreased and black: undetected. Genes are depicted by circles, red: increased, blue: decreased. Solid arrows indicate direct reaction, dotted arrows indicate the reaction is broken by one or more sub-reactions. The tissue in which genes or metabolites were significantly altered are depicted by the presence of a muscle, spinal cord or liver image. The timepoint at which a gene or metabolite was detected as significantly changed is indicated in months (M). Image constructed using Biorender.

**Supplementary Figure 25.**
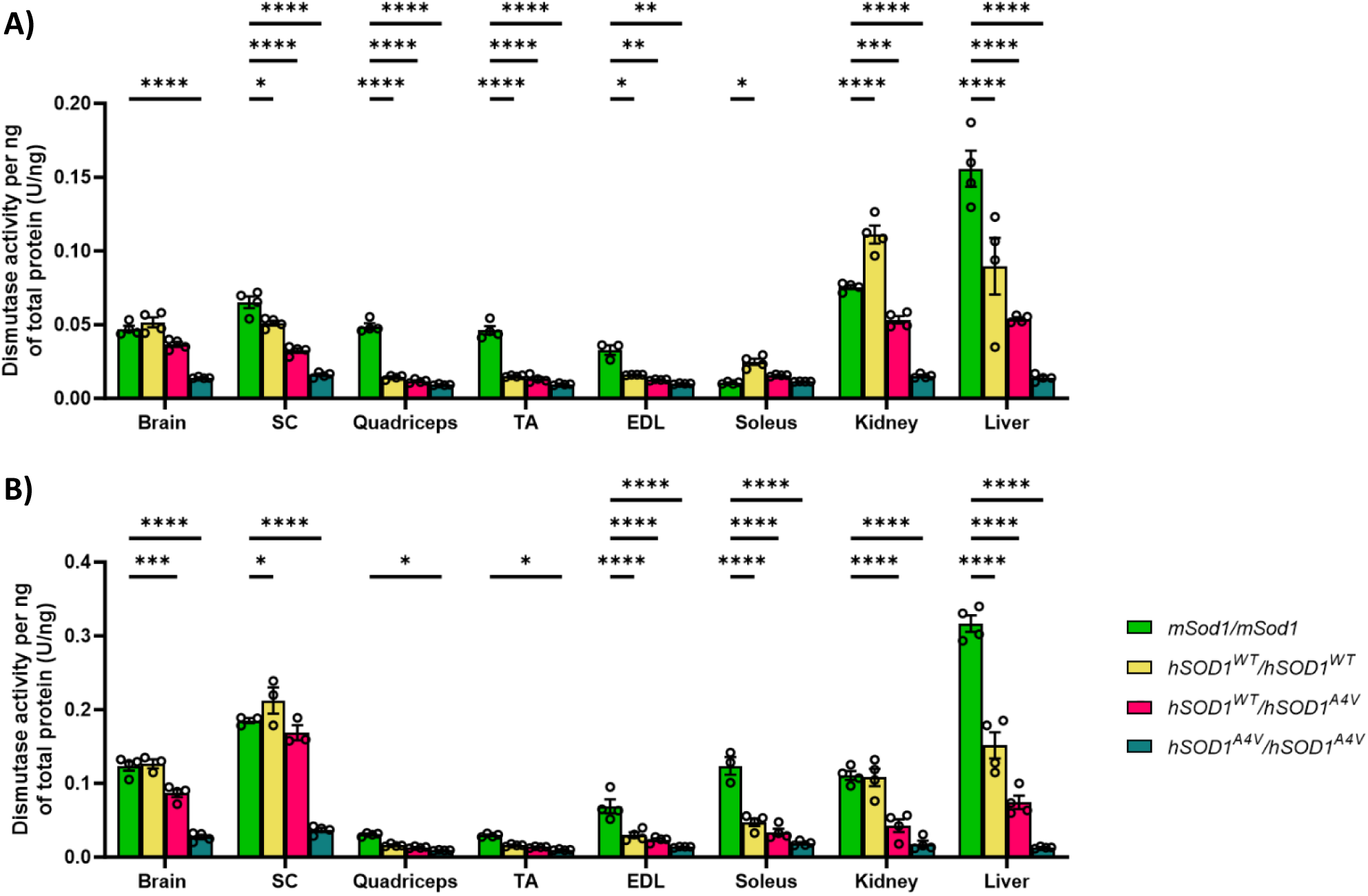
Effect of SOD1 humanisation on SOD1 activity. Spectrophotometric dismutase activity assays in tissues from A) male and B) female *hSOD1^A4V^*mice and 4-month-old *mSOD1/mSOD1* controls. Mean +/- SEM. One-way ANOVA with Tukey’s post hoc test. For clarity, only significant comparisons to *mSOD1* controls are shown. * p < 0.05, ** p < 0.01, *** p < 0.001, **** p < 0.00001. ns, not significant; SC, spinal cord; TA, tibialis anterior; EDL, extensor digitorium longus.

